# Distinct clades of TELOMERE REPEAT BINDING transcriptional regulators interplay to regulate plant development

**DOI:** 10.1101/2023.08.16.553498

**Authors:** Simon Amiard, Léa Feit, Lauriane Simon, Samuel Le Goff, Loriane Loizeau, Léa Wolff, Falk Butter, Clara Bourbousse, Fredy Barneche, Christophe Tatout, Aline. V. Probst

## Abstract

TELOMERE REPEAT BINDING proteins (TRBs) are plant-specific transcriptional regulators that combine two DNA-binding domains, the GH1 domain shared with H1 histones that binds to linker DNA and the Myb/SANT domain that specifically recognizes the telobox DNA binding site motif. TRB1, TRB2 and TRB3 proteins recruit the Polycomb group complex 2 (PRC2) to deposit H3K27me3 and JMJ14 to remove H3K4me3 at target genes containing telobox motifs in their promoters to repress transcription. Here, we characterize the function of TRB4 and TRB5, which belong to a separate TRB clade conserved in spermatophytes. TRB4 and TRB5 affect the transcriptional control of several hundred genes involved in developmental responses to environmental cues, the majority of which differ from differentially regulated genes in *trb1 trb2 trb3,* suggesting distinct modes of action at the chromatin level. Indeed, TRB4 binds to several thousand sites in the genome, mainly at TSS and promoter regions of transcriptionally active and H3K4me3-marked genes but is not enriched at H3K27me3-marked gene bodies. TRB4 physically interacts with the PRC2 component CURLY LEAF (CLF), but, unexpectedly, loss of TRB4 and TRB5 partially suppresses the developmental defects of *clf* mutant plants, by acting as transcriptional activators of the key flowering genes *SOC1* and *FT.* We further show that TRB4 and TRB1 share multiple target genes and reveal physical and genetic interactions between TRBs of the two distinct clades, collectively unveiling that TRB proteins engage in both positive and negative interactions with other members of the family to regulate plant development through PRC2-dependent and independent mechanisms.

## Introduction

Development and adequate response to the environment requires sophisticated mechanisms to precisely regulate gene expression. Conserved through evolution, the Polycomb group (Margueron and Reinberg, 2011; Schuettengruber et al., 2017) (PcG) proteins restrict gene activity during development by placing repressive histone modifications, while proteins from the Trithorax group (TrxG) set histone modifications permissive for transcription and can thereby counteract the PcG machinery. PcG function is essential for development, as loss of Polycomb activity is lethal in mice (Sauvageau and Sauvageau, 2008), causes ectopic expression of Homeotic (*Hox*) genes in Drosophila (Lewis, 1978)and induces severe developmental phenotypes in Arabidopsis (Simmons and Bergmann, 2016).

In animals and plants, Polycomb repressive complexes (PRC) can be classified by their distinct enzymatic activities. While PRC1 comprises histone H2A ubiquitination activity, the PRC2 complexes harbor a histone methyltransferase that trimethylates lysine 27 of histone H3 (H3K27me3). In Drosophila, the PRC2 complex consists of four subunits: the catalytic subunit Enhancer of zeste (E(z)); Extra sex combs (Esc); Suppressor of zeste 12 (Su(z)12), critical for nucleosome binding; and Nuclear remodeling factor 55 (NURF55).

In Arabidopsis, PRC2 complexes have undergone functional diversification with three catalytic SET-domain proteins namely CURLY LEAF (CLF) and SWINGER (SWN) that are responsible for the sporophytic H3K27me3 activity and MEDEA (MEA) that functions in repressing endosperm proliferation (Vijayanathan et al., 2022) as well as three Su(z)12 orthologs termed VERNALIZATION 2 (VRN2), EMBRYONIC FLOWER 2 (EMF2) and FERTILIZATION INDEPENDENT SEED 2 (FIS2). This enlargement of the E(z) and Su(z)12 gene families allows assembly of several different PRC2 complexes that operate at diverse developmental stages ensuring important developmental transitions (Wang et al., 2016). Single mutants lacking only one of the possible catalytic subunits of PRC2 are viable in Arabidopsis but show severe developmental abnormalities (Goodrich et al., 1997). Genome-wide profiling in Arabidopsis identified 7000 to 8000 thousand genes enriched in the H3K27me3 mark, *i.e*., approximately one third of all protein-coding genes. These genes are in general weakly expressed, involved in development, but also in response to stress (Zhang et al., 2007).

PRC2 complexes need to be recruited to different target genes in a sequence-specific manner. PRC2 core subunits do not harbor intrinsic DNA binding activity, and recruitment to target genes therefore involves associated proteins that, despite the evolutionary conservation of the PRC2 core components, differ widely between species. In Drosophila, PRC2 is recruited via DNA-binding proteins at Polycomb Response Elements (PREs) situated in regulatory regions of genes (Horard et al., 2000). In mammals, hypomethylated CpG islands may represent PRE-like sequences, to which PRC2 can be targeted via interaction with several Transcription Factors (TFs) or via non-coding RNAs (Davidovich and Cech, 2015). Binding of the Esc homolog EED to H3K27me3 methylated histones further stabilizes PRC2 and stimulates the histone methyltransferase activity of the complex (Margueron et al., 2009). PRC2 recruitment via long non-coding RNA has also been evidenced in plants (Ariel et al., 2014), and several *cis*-elements with PRE-like characteristics have been identified and linked to PcG recruitment in Arabidopsis. These include a six-nucleotide RY motif (Yuan et al., 2020), RLE element in the *LEAFY COTYLEDON 2 (LEC2)* gene promoter (Berger et al., 2011), the ASYMMETRIC LEAVES1 (AS1) and AS2 binding sites in *BREVIPEDICELLUS* and *KNAT2* promoters (Lodha et al., 2013) or GAGA and telobox motifs (Deng et al., 2013; Zhou et al., 2016; Xiao et al., 2017), short motifs abundant in gene promoters that in turn are bound by specific proteins recruiting the Polycomb complexes.

A group of such plant-specific proteins that bind preferentially to telomeric motifs (so-called ‘teloboxes’) via their Myb-like DNA binding domain is the TELOMERE REPEAT BINDING (TRB) protein family consisting of five members (TRB1 - 5) in Arabidopsis (Schrumpfov et al., 2004; Schrumpfová et al., 2014). TRB1, TRB2 and TRB3 were initially identified to bind telomeres consisting of long tandem repeats of teloboxes and were proposed to function in telomere protection (Schrumpfov et al., 2004; Mozgová et al., 2008). Besides the N-terminal Myb-domain, TRB proteins comprise a second DNA binding domain, a globular H1 (GH1) domain, shared with linker histones H1, where it mediates binding to the nucleosome dyad and linker DNA (Bednar et al., 2017). The GH1 domain is also involved in TRB protein-protein interactions including TRB1 homodimerization(Mozgová et al., 2008) and its heterodimerization with TRB2 and TRB3 (Schrumpfová et al., 2008). TRB1-3 proteins finally comprise a coiled-coil region in their C-termini that interacts with the catalytic PRC2 subunits CLF and SWN, triggering H3K27me3 deposition at a subset of PRC2 target genes (Zhou et al., 2016; Zhou et al., 2018). This role in gene repression is further reinforced by the interaction of TRB1, TRB2 and TRB3 with the JMJ14 H3K4me3 demethylase thereby both counteracting the maintenance of a transcriptionally permissive and establishing a repressive chromatin state (Wang et al., 2023). TRB1-3 mediated recruitment of PRC2 activity is restricted by specific chromatin characteristics such as the presence of linker histone H1, as in absence of H1, TRB1 accumulates at telomeres and at Interstitial Telomere Repeat sequences (ITRs) within the pericentromeric regions of specific chromosomes leading to H3K27me3 accumulation at these sequences (Teano et al., 2022). In addition to their function in PRC2 targeting, TRB1 has also been shown to maintain high expression levels of genes involved in metabolism such as photosynthesis (Zhou et al., 2016) revealing a yet poorly understood mode of action that may depend on target genes, other transcriptional regulators and chromatin context. Finally, TRB1 and TRB2 are also members of the PEAT (PWWPs–EPCRs–ARIDs–TRBs) complex, which is required for histone deacetylation at transposable elements (TEs) and heterochromatin silencing (Tan et al., 2018).

In Arabidopsis, TRB1, TRB2 and TRB3 seem to fulfill redundant roles, as phenotypes of neither single nor double mutants are distinguishable from those of wild type plants. Triple *trb1 trb2 trb3* mutants, however, harbor strong developmental defects like those observed in mutants lacking PRC2 complex activity (Zhou et al., 2018). While several functions of TRB1-3 have been characterized, it is yet unclear whether and how other TRB family members contribute to gene expression regulation and plant development.

Here, we show that the TRB family separated in two clades comprising either Brassicaceae TRB1-3 or TRB4-5 proteins at the moment of the appearance of seed plants. We find that Arabidopsis TRB4 and TRB5 are nuclear proteins that act redundantly in developmental regulation. Genome-wide profiling identified TRB4 to preferentially associate with promoter sequences of transcriptionally active genes, and showed that, in agreement with our observation that all TRB proteins engage in homo- and heterodimerization, TRB1 and TRB4 target loci substantially overlap. However, while TRB1 accumulates at gene bodies of a subset of H3K27me3-enriched genes, TRB4 accumulates at transcription start and promoter regions of H3K4me3-enriched and highly transcriptionally active genes. Genes misregulated in *trb4 trb5* double mutant plants are overrepresented among genes showing co-occurrence of H3K4me3 and H3K27me3, yet the loss of TRB4 and TRB5 only affects enrichment of these histone modifications at a small subset of genes suggesting that they act independently of these histone marks. Finally, we reveal that TRB4 and TRB5 physically and genetically interact with PRC2 subunits and are unexpectedly required for the early flowering and leaf curling phenotype of mutants lacking the catalytic subunit CLF likely by acting as positive transcriptional regulators of *FT* and *SOC1*. We suggest that TRB4 and TRB5 proteins function in PcG-dependent and independent gene regulation to fine-tune gene expression during development in concert with the other members of the TRB family.

## Results

### TRB4 and TRB5 proteins belong to a separate phylogenetic clade

In an initial attempt to identify proteins with the capacity to bind telomere repeats, we performed a label-free quantitative proteomics analysis of proteins binding to the Arabidopsis TTTAGGG repeat sequence(Charbonnel et al., 2018). In addition to TRB1, TRB2 and TRB3, data re-analysis identified two poorly characterized members of the TRB family, namely TRB4 and TRB5, which are significantly enriched (Fold Change (FC) of 2 and 7.3 respectively) in the telomere pull-down compared to the shuffled DNA control (***Figure 1A***). Previous studies suggested that TRB4 and TRB5 belong to a separate clade in the TRB phylogeny (Kotlinski et al., 2017; Kusová et al., 2023). To time the appearance of this clade, we selected 24 species to represent the diversity of the green land plant lineage and interrogated several databases using Arabidopsis TRB1 to TRB5 as query. Except for unicellular algae where TRB orthologs were not found, a parental TRB protein containing both an amino-terminal Myb/SANT domain and a central GH1 domain is present in bryophytes and was subject to duplication and diversification in an ancestral species of spermatophytes (***Figure 1B, Supplementary Table 1***). Phylogenetic analysis using IQ-Tree (***Supplementary Figure 1A***) identified two separate TRB clades conserved in both gymnosperms and angiosperms that comprise either Arabidopsis TRB1-3 or TRB4-5, which we termed clade I and clade II respectively. Longer branch lengths were observed for clade II compared to clade I (0.27 versus 0.39 substitutions per site per species in clade I and clade II respectively, p < 0,0062) indicating higher evolutionary divergence in clade II. Within each clade, TRB proteins of gymnosperms, monocotyledons and dicotyledons respectively group together. In dicotyledons, *TRB* genes underwent further expansion and sub-functionalization such as within clade I, where TRB proteins further diverged from a common ancestor into a TRB1 and a TRB2/3 sub clade. We also noticed that TRB1 and TRB4/5 orthologs are present in all dicotyledons, while representatives of the TRB2/3 sub-clade are absent in certain species (***Figure 1B***).

**Figure 1:**
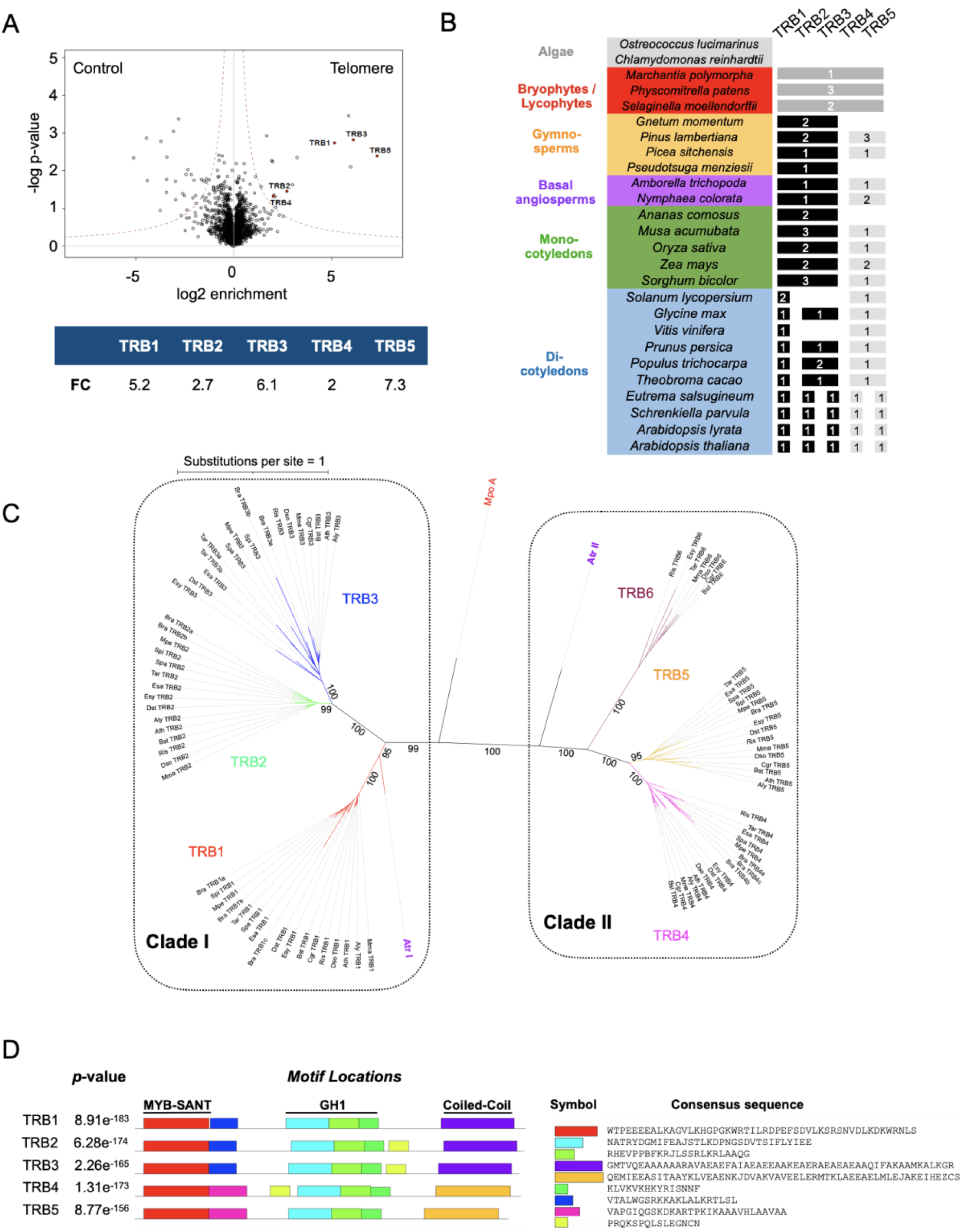
TRB4 and TRB5 bind to telomere repeats and belong to a separate TRB clade conserved in gymnosperms and angiosperms. **(A)** Volcano plot showing enrichment of the five TRB proteins (red dots) in the telomere repeat pull-down relative to scrambled control sequences. Mean enrichment values from 4 independent experiments are indicated. **(B)** Presence or absence of orthologs of the Arabidopsis TRB proteins in different plant species spanning the evolutionary history of land plants. TRB orthologs can be detected in the liverwort Marchantia and orthologs of TRB1-3 (black) and TRB4-5 (grey) exist in gymnosperms, basal angiosperms, mono- and di-cotyledons. **(C)** Unrooted maximum likelihood phylogenetic tree of 87 TRB orthologs from 15 Brassicaceae species. **(D)** Presentation of 9 motifs in the *Arabidopsis thaliana* TRB proteins predicted with MEME from the alignment of the TRB orthologs from the 15 Brassicaceae species. Sequences of consensus motifs are indicated. Distinct motifs adjacent to the MYB/SANT domain and different coiled-coil domains in the C-termini differentiate clade I from clade II TRB proteins.

Considering that TRB proteins have diversified even further within the Brassicaceae, we analyzed in more detail the phylogenetic relationship of 87 TRB orthologs within 15 species of this plant family (***Figure 1C***). The non-rooted phylogenetic tree confirms that within clade I, the TRB1 and the TRB2/3 subclades resulted from duplication of a common ancestor gene and that TRB2 and TRB3 on the one hand, and TRB4 and TRB5 on the other, diverged after more recent duplications from ancestors within each subclade (***Figure 1C***). Finally, 7 out of the 15 Brassicaceae species analyzed encode proteins grouping into an additional TRB subclade belonging to clade II TRBs that we termed TRB6 (***Figure 1C***).

Using all TRB orthologs from the Brassicaceae family as input, we then predicted protein motifs (***Figure 1D, Supplementary Figure 1B***). MEME reveals motifs specific to each clade adjacent to the Myb/SANT domain and in the C-terminal part of the TRB proteins (***Figure 1D***). The latter are identified by InterProScan as coiled-coil domains and are predicted by alpha-fold (Jumper et al., 2021) to form long alpha helices (***Supplementary Figure 1C***). Most TRB1 proteins further comprise a supplementary motif specific to TRB1 orthologs (***Supplementary Figure 1B***). Finally, a short motif present between the GH1 and the coiled-coil domain in most clade I TRB proteins (***Supplementary Figure 1B***) is also found in TRB4, although at a different position within the protein suggesting genomic rearrangements.

In summary, TRB4 and TRB5 share the two DNA binding domains with TRB1-3. Nevertheless, TRB proteins clearly separate into two phylogenetic clades distinguished by divergent regions, in particular the coil-coiled C-terminal region, involved in protein-protein interaction between TRB1-3 and the PRC2 complex (Zhou et al., 2018), thus opening the perspective of functional diversification of clade II TRB proteins.

### TRB4 and TRB5 fine tune plant development and gene expression, but are dispensable for telomere protection

Given their interaction with telomere repeats, we first investigated whether TRB4 and TRB5 play a role in telomere regulation or stability. We generated CRISPR-Cas9 loss-of-function alleles by targeting Cas9 to the first exon of *TRB4* and the second exon of *TRB5*. For each gene we retained two independent mutant alleles, in which nucleotide insertions or deletions led to frame shifts resulting in premature stop codons (***Supplementary Figure 2A***). All mutants are therefore expected to be null mutants. *TRB4* or *TRB5* loss-of function plants did not show any developmental abnormalities **(Figure 2A)** or alterations in telomere maintenance, as determined by quantifying the number of anaphase bridges in inflorescences and γH2A.X foci in root tip nuclei **(Figure 2B)** or by testing potential telomere de-protection using Telomere Restriction Fragments (TRF) analysis of bulk telomere length (**Figure 2C)**. Considering that *TRB4* and *TRB5* may be functionally redundant, we generated *trb4 trb5* double mutants that also showed no defects in telomere maintenance (**Figure 2B-C**). Hence, while these two proteins target telomeric DNA, their removal is not sufficient for telomere deprotection.

**Figure 2:**
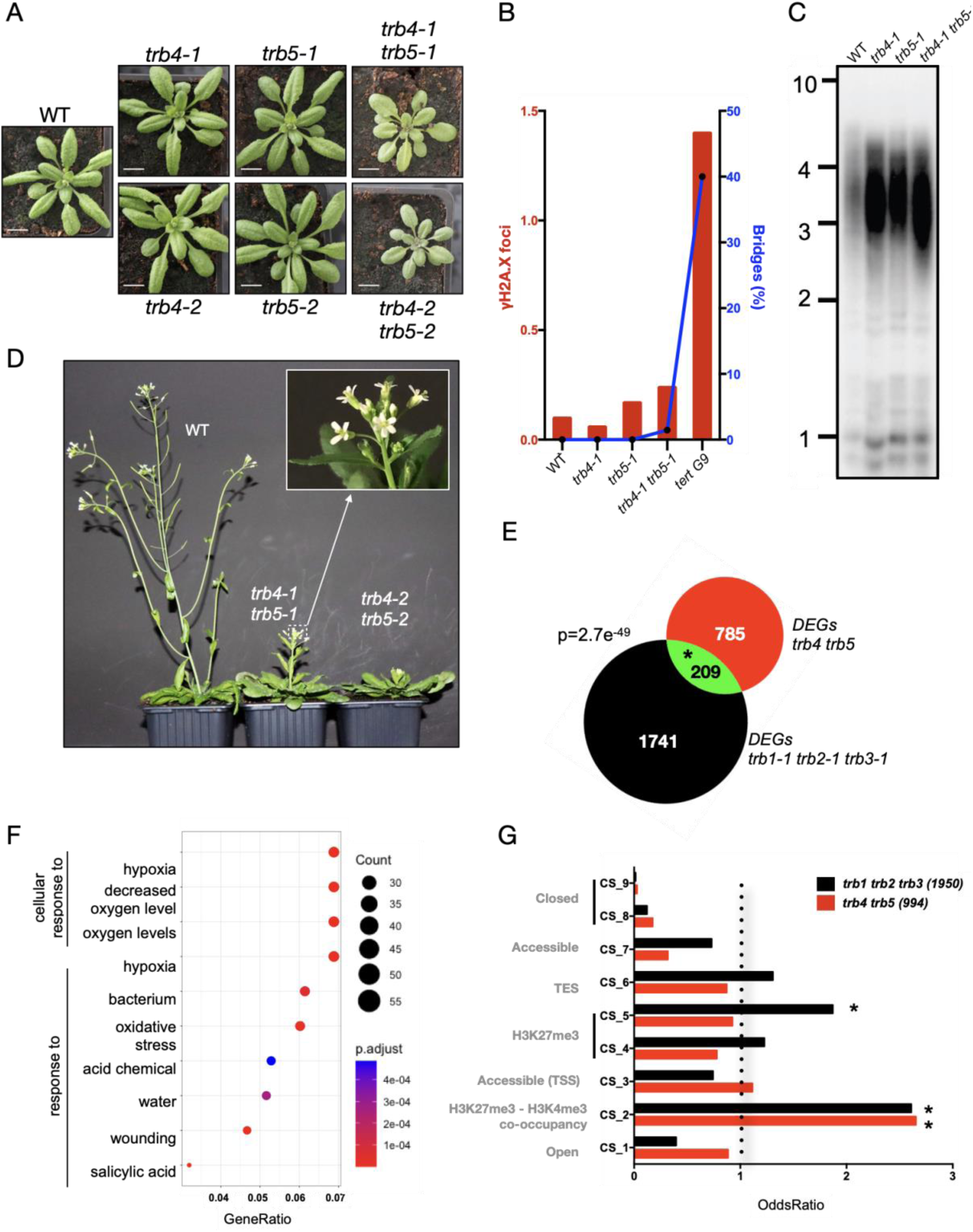
TRB4 and TRB5 are transcriptional regulators required for plant development, but do not for telomere protection. **(A)** Representative wild type, *trb4-1, trb5-1, trb4-2, trb5-2 single and* double mutant plants at three weeks of age. **(B)** Percentage of anaphase bridges (blue line) and mean number of γH2A.X foci (histogram, red) in wild type, *trb4-1* and *trb5-1* single and *trb4-1 trb5-1* double mutants. Plants lacking the TElomerase Reverse Transcriptase TERT in the 9^th^ generation (*tert G9*) (Fitzgerald et al., 1999) were used as positive control of telomere deprotection. **(C)** Telomere Restriction Fragments (TRF) analysis of bulk telomere length in genomic DNA using telomere repeat probes in wild type, *trb4-1* and *trb5-1* single and *trb4-1 trb5-1* double mutants. **(D)** Representative single and double mutant plants at the flowering stage. Double mutants show delayed flowering and supernumerary petals (quantification in ***Supplementary Figure 2E, F***). **(E)** Venn diagram showing number of Differentially Expressed Genes (DEGs) in *trb4 trb5* and *trb1 trb2 trb3* mutants and those common to both mutant combinations. Significance of common DEGs was determined using a hypergeometric test. **(F)** Gene Ontology (GO) term enrichment of *trb4 trb5* DEGs defined using ClusterProfile. **(G)** Enrichment of *trb4 trb5* and *trb1 trb2 trb3* DEGs in the nine chromatin states defined by (Sequeira-Mendes et al., 2014) (*OR>1 and p<0.05).

However, we noticed several important developmental abnormalities in *trb4 trb5* double mutants: young seedlings are smaller compared to wild type plants or single mutants, show brighter leaf color and shorter roots (***Figure 2A, Supplementary Figure 2B)***. Adult plants display delayed flowering and reduced fertility (***Figure 2D; Supplementary Figure 2C, 2E***). Furthermore, ∼30% of *trb4-1 trb5-1* double mutant flowers harbor supernumerary petals **(Figure 2D (inset)*, Supplementary Figure 2F)***, a phenotype previously observed in mutants for histone H3K27 demethylases(Yan et al., 2018) and for TrxG mutants (Carles et al., 2005). Hence, complementation of *trb4-1 trb5-1* mutant with either *TRB4* or *TRB5* expressed under their respective endogenous promoter led to a fully restored WT phenotype confirming that TRB4 and TRB5 harbor redundant functions (***Supplementary Figure 2D, E, F***).

To address how TRB4 and TRB5 impact plant development, we carried out RNA-seq in 7-day-old seedlings of WT, the two independent *trb4 trb5* double mutant lines and a *trb1-1 trb2-1 trb3-1* triple mutant line. We retained 994 differentially expressed genes (DEGs), 62% being upregulated and 38% downregulated, shared between mutant plants combining distinct *trb4 trb5* mutant alleles (***Supplementary Figure 2G***). More than half of the misregulated genes are categorized to function in ‘response to stress’ and are linked to the cellular response to hypoxia, oxygen, light, and hormone levels (***Figure 2F***) implying that the plant’s response to environmental stimuli is affected upon loss of TRB4 and TRB5. Several genes encoding transcription factors belonging to the AP2/ERF, homeobox and MADS-box transcription factor families are also misregulated including the flowering regulator genes *SUPPRESSOR OF OVEREXPRESSION OF CO 1 (SOC1)* and *FLOWERING LOCUS C (FLC)*.

The number of DEGs in *trb4 trb5* plants is lower (n=994) than the number of DEGs in *trb1-1 trb2-1 trb3-1* triple mutants (n=1950) shared among our dataset and a previous one (Zhou et al., 2018) (***Figure 2E, Supplementary Figure 2H***), in agreement with the milder developmental phenotype of these plant lines. Yet, *trb4 trb5* plants share 21% of their DEGs with *trb1 trb2 trb3* triple mutants both oppositely and co-regulated (***Supplementary Figure 2I***), showing that subsets of genes are directly or indirectly regulated by members of both TRB clades. To explore whether loss of TRB4 and TRB5 preferentially affects genes with a particular chromatin state (CS), we analyzed whether the TSS of the misregulated genes are characterized by any of the previously identified CS (Sequeira-Mendes et al., 2014) (***Figure 2G***). In agreement with the involvement of TRB1-3 in PcG-mediated transcriptional control, genes differentially regulated in *trb1 trb2 trb3* are overrepresented among genes corresponding to CS5 (H3K27me3-rich) and genes showing co-occurrence of both H3K4me3 and H3K27me3 modifications (CS2) (Sequeira-Mendes et al., 2014). Instead, *trb4 trb5* DEGs are mainly overrepresented among genes associated with CS2 but not with CS5. While TRB1-3 proteins are involved in TE silencing as part of the PEAT complex (Tan et al., 2018), we did not observe a strong reactivation of TEs in *trb4 trb5* mutant plants (17 TEs up, 13 TEs down), altogether showing that a major function of TRB4-5 proteins is gene expression control.

### TRB proteins of the two clades physically and genetically interact with each other

Clade I TRB proteins were previously shown to interact with each other through their GH1 domain (Schrumpfová et al., 2008). We therefore envisaged that similar interactions could take place within clade II (TRB4-5) or between the two clades. To test this, we carried out yeast-two-hybrid (Y2H) experiments using each protein either as bait or as prey. Our assay confirmed known interactions among clade I and, in agreement with (Kusová et al., 2023), showed that TRB4 and TRB5 can homo- and heterodimerize in yeast (***Figure 3A, Supplementary Figure 3A***). We further observed that TRB4 interacts with TRB2 and TRB3, indicating that interactions between TRB proteins of different clades can take place. To test the occurrence of these interactions *in planta*, we used Bimolecular Fluorescence Complementation (BiFC) assays in *N. benthamiana* leaves. BiFC confirmed the protein-protein interactions identified using Y2H and revealed additional interactions between TRB1 and TRB4, as well as between TRB5 and TRB1-3 **(*Figure 3B*).** Protein-protein interactions between the different TRB clades all take place in the nucleus and concentrate in few bright nuclear speckles likely corresponding to *N. benthamiana* telomeres as observed in (Schrumpfová et al., 2014).

**Figure 3:**
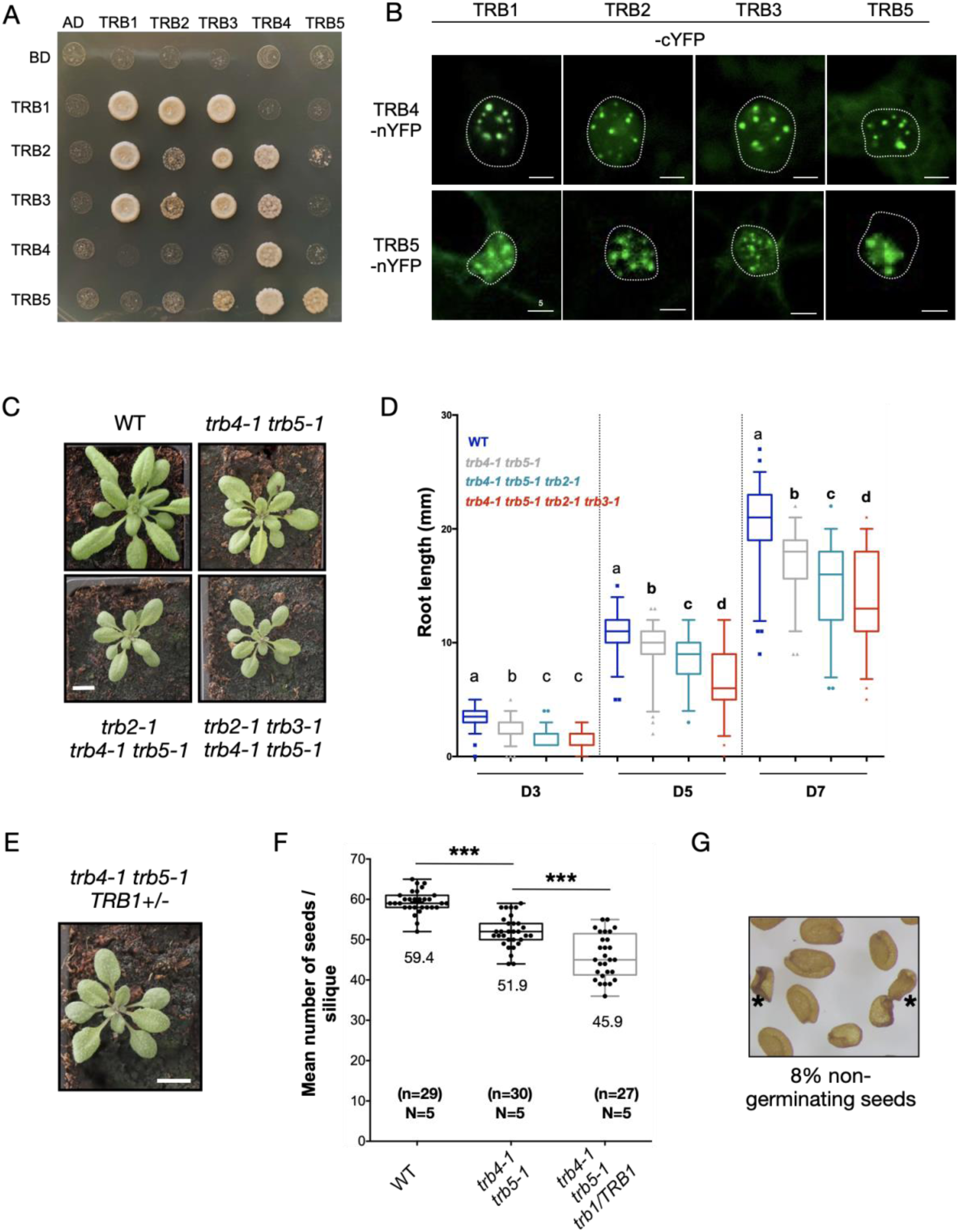
TRB proteins from the two clades physically and genetically interact with each other. **(A)** Interaction of the five Arabidopsis TRB proteins with each other probed in the Yeast-Two-Hybrid system. Growth on selective medium lacking histidine and adenine reveals interaction between the two proteins tested. Horizontal: Translational fusions with the Gal4-Activation domain (AD). Vertical: Translational fusion with the Gal4-DNA Binding Domain (BD). BD and AD indicate the respective empty vectors. **(B)** Bimolecular Fluorescence Complementation reveals protein-protein interactions (PPIs) between TRB proteins within each clade and between the proteins belonging to the TRB_I and TRB_II clade in *N. benthaniama* leaf cells. Maximum intensity projections of Z-stacks acquired with a confocal microscope are shown. PPI takes place within distinct nuclear speckles likely corresponding to telomeres. **(C)** Representative wild type, *trb4-1 trb5-1* double, *trb2-1 trb4-1 trb5-1* triple and *trb2-1 trb3-1 trb4-1 trb5-1* quadruple mutant plants at three weeks of age. Scale bar = 1cm. **(D)** Quantification of root length from *in vitro* grown wild type, *trb4-1 trb5-1* double, *trb2-1 trb4-1 trb5-1* triple and *trb2-1 trb3-1 trb4-1 trb5-1* quadruple mutants at day 3 (D3), 5 and 7 after germination. For each time point, values from two independent replicates are shown. Different letters indicate significant differences among samples of the same time point by Mann-Whitney test (p < 0.01). **(E)** Representative *trb4-1 trb5-1 trb1-1*/*TRB1* mutant plant. Scale bar = 1cm. **(F)** Mean number of seeds per silique from wild type, *trb4-1 trb5-1*, and *trb4-1 trb5-1 trb1-1*/*TRB1* plants. At least 27 siliques from five plants were counted. About 12 % less seeds are present in the *trb4-1 trb5-1* mother plant heterozygous for the *trb1* mutation compared to the *trb4-1 trb5-1* double mutants (***p < 0.0001, t-test). **(G)** Seeds from *trb4-1 trb5-1 trb1-1*/*TRB1* plants revealing the presence of shriveled, non-germinating seeds marked with an asterisk.

Given the physical interaction between the different TRB proteins, we explored their genetic interaction in multiple mutants obtained by crossing first *trb1-1, trb2-1* or *trb3-1* single mutants with *trb4-1 trb5-1* plants and subsequent crosses of the resulting multiple mutants. From the segregating populations, we obtained viable *trb2-1 trb4-1 trb5-1* triple and *trb2-1 trb3-1 trb4-1 trb5-1* quadruple mutant plants, which strongly resemble clade II *trb* mutants but show more pronounced developmental deficiencies including smaller rosettes (***Figure 3C***) and aggravated root growth defects (***Figure 3D***).

In contrast, all attempts to obtain *trb1-1 trb4-1 trb5-1* triple mutants failed. Closer inspection of the siliques from *trb4-1 trb5-1 trb1-1/TRB1* plants (***Figure 3E***) revealed aborted ovules and a smaller number of seeds per silique (***Figure 3F***), indicating failed fertilization or early abortion of the developing seed. However, less than 25% of the ovules abort suggesting that a fraction of triple mutants completes seed development. Indeed, we find that 8% (41/509) of the seeds fail to germinate (***Figure 3G***), so that all surviving plantlets are either WT or heterozygous for the *trb1* mutation (65% *TRB1/trb1* and 35% *TRB1/TRB1*, n = 89). In absence of clade II TRB proteins, TRB1 therefore fulfills an essential function that cannot be complemented by TRB2 or TRB3. The requirement for TRB1 in the *trb4 trb5* mutant background might be explained by its higher expression in embryo and endosperm (***Supplementary Figure 3C***) or by a yet undefined specific role of TRB1, the only clade I TRB protein present in all dicotyledonous species analyzed (***Figure 1B, left***).

### TRB4 binds preferentially to promoter regions

To study the localization of TRB4 and TRB5, we expressed both proteins as a translational fusion with GFP. As estimated from a full restoration of wild type flowering time and normal petal development in the *trb4 trb5* double mutant background, these TRB4-GFP and TRB5-GFP fusion proteins are functional (***Figure 4A, B***). We first imaged GFP fluorescence in root tips of young plantlets and found TRB4 and TRB5 to localize to the nucleus (***Figure 4C***) as previously observed for TRB1, TRB2 and TRB3 (Schrumpfová et al., 2014; Zhou et al., 2018) and confirmed here for TRB1 (***Figure 4C***). Making use of these transgenic plants, we carried out immunofluorescence staining in isolated nuclei from 7-day-old seedlings, to examine the subnuclear localization of TRB4, TRB5 and TRB1 in more detail. The three proteins localize throughout euchromatin, sometimes as small speckles, but are depleted from the DAPI-bright heterochromatic chromocenters (***Figure 4D***). TRB4 and TRB5 as well as TRB1 are also detected in the nucleolus as was already observed after transient expression of TRBs in tobacco leaves (Zhou et al., 2016; Kusová et al., 2023). To obtain a precise view on the genomic distribution of a clade II TRB proteins, we carried out ChIP-seq targeting TRB4-GFP in 7-day old plantlets and identified more than 5000 TRB4 peaks robustly detected in two independent biological replicates (***Supplementary Figure 4A***). In agreement with our microscopic observations, TRB4-GFP associated loci are enriched at chromosome arms and depleted from pericentromeric heterochromatic regions (***Figure 4E***). Over 68% of the TRB4 peaks are situated in gene promoters (***Figure 4F***). *De novo* motif discovery identified like for TRB1 (Schrumpfová et al., 2015; Zhou et al., 2016; Teano et al., 2022) the ‘telobox’ consensus motif (*TAGGGTT*) as the most enriched motif present at about 49% of TRB4 genomic binding sites (MEME, p=7.4^e-32^). TRB4-GFP was also significantly enriched at loci bearing the ‘site II motif’ (*TGGGCY*) typically associated with the telobox motif in promoters of ribosomal genes (Gaspin et al., 2010) (***Supplementary Figure 4B***). Therefore, TRB4 binds preferentially to promoters and TSSs, many of which carry telobox motifs, although TRB4 is also present at telobox-free sites through a recruitment mode that remains to be discovered.

**Figure 4:**
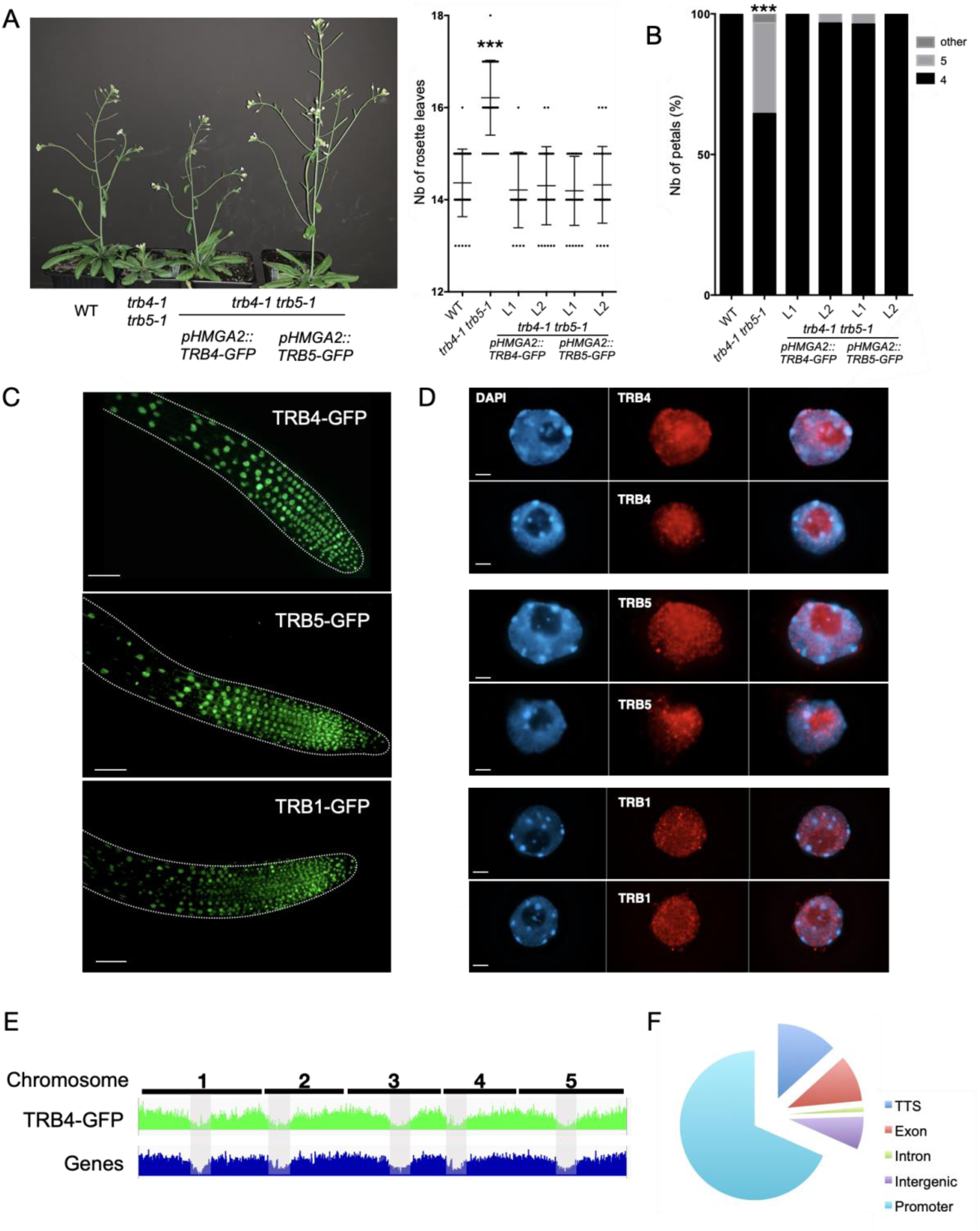
TRB4 and TRB5 are nuclear proteins enriched in euchromatin and TRB4 preferentially binds to gene promoters. TRB4-GFP or TRB5-GFP fusion proteins expressed under the *GH1-HMGA2* promoter complement the late flowering and supernumerary petal phenotype of *trb4-1 trb5-1* double mutants. **(A)** Representative 4 weeks old plants (left). Quantification (right) of the number of leaves at bolting in wild type, *trb4-1 trb5-1* double mutants and four independent transgenic lines expressing TRB4-GFP or TRB5-GFP. *** p < 0.0001, t-test. **(B)** Percentage of flowers showing 4, 5 or any other aberrant petal number in the same genotypes as in (**A**). *** p<0.001, t-test. **(C)** Representative root tips of plants expressing TRB4, TRB5 or TRB1 as GFP fusions. Fusion proteins localize in the nucleus. Scale bar corresponds to 50 μm. **(D)** Maximum intensity projections of nuclei from plantlets expressing TRB4, TRB5 or TRB1 as GFP fusions, in which fusion proteins were revealed with an anti-GFP antibody (red) by immunofluorescence staining. TRB4, TRB5 and TRB1 localize to small discrete speckles throughout euchromatin. DNA is counterstained with DAPI (blue, left). Merged images are shown on the right. Scale bar corresponds to 2 μm. **(E)** Comparison of the distribution of the TRB4-GFP ChIP-seq peaks and genes along the 5 Arabidopsis chromosomes. Grey zones indicate centromeric and pericentromeric regions. (**F**) Distribution of TRB4 peaks determined by ChIP-seq among different genomic features in the Arabidopsis genome.

### TRB4 and TRB1 show different binding patterns along genes but share targets and engage in a complex gene co-regulation

To obtain a detailed view on TRB1 and TRB4 differential binding, we carried out k-mean clustering of their target genes (**Figure 5A**). Within cluster 1 and 2, TRB4 strongly marks the TSS while cluster 3 contains genes showing TRB4 binding further upstream and/or downstream of the TSS. Motif analyses of the 5’ UTR region of cluster 1 and 2 genes indicate a strong enrichment (E-value > 1^e-200^) in telobox motifs, while no significant enrichment of this motif was found in the promotor and 5’UTR regions (-1000bp) of genes from cluster 3 (***Supplementary Figure 5***). Genes in cluster 1 are significantly more expressed than the average of all TRB4 target genes or those in cluster 2 and 3 (***Figure 5D)***. Similar to TRB4, k-mean clustering of TRB1 binding sites identified in our recent study (Teano et al., 2022) distinguished a group with enrichment at the TSS (TRB1 cluster 2) similar to TRB4 cluster 1 and 2 (**Figure 5B**). TRB1 cluster 1 on the contrary corresponds to a group of genes, for which TRB1 marks the entire gene body and for which no ‘Telobox’ motif enrichment was found either in the 5’UTR nor in the coding sequences (***Supplementary Figure 5***). Plotting TRB1 and TRB4 on TRB1 cluster 1 genes indicates that, except for a few genes, they are preferentially enriched in TRB1 but not TRB4 (***Figure 5C)***. Interestingly, genes targeted by TRB4 or TRB1 at their TSS (TRB4 cluster1 and 2, TRB1 cluster2) are frequently involved in ribosomes biogenesis and translation, in agreement with the enrichment of telobox and site II motifs in their promoters, whereas genes fully covered by TRB1 (TRB1 cluster 1) are frequently involved in stress and developmental responses (***Supplementary Figure 5***). Plotting mean gene expression levels for each cluster identifies that TRB1 specifically targets a group of lowly expressed genes (cluster 1) while in contrast TRB4 targets a group of genes that are particularly highly expressed (***Figure 5D***), illustrating a specialization of the members from the different TRB clades.

**Figure 5:**
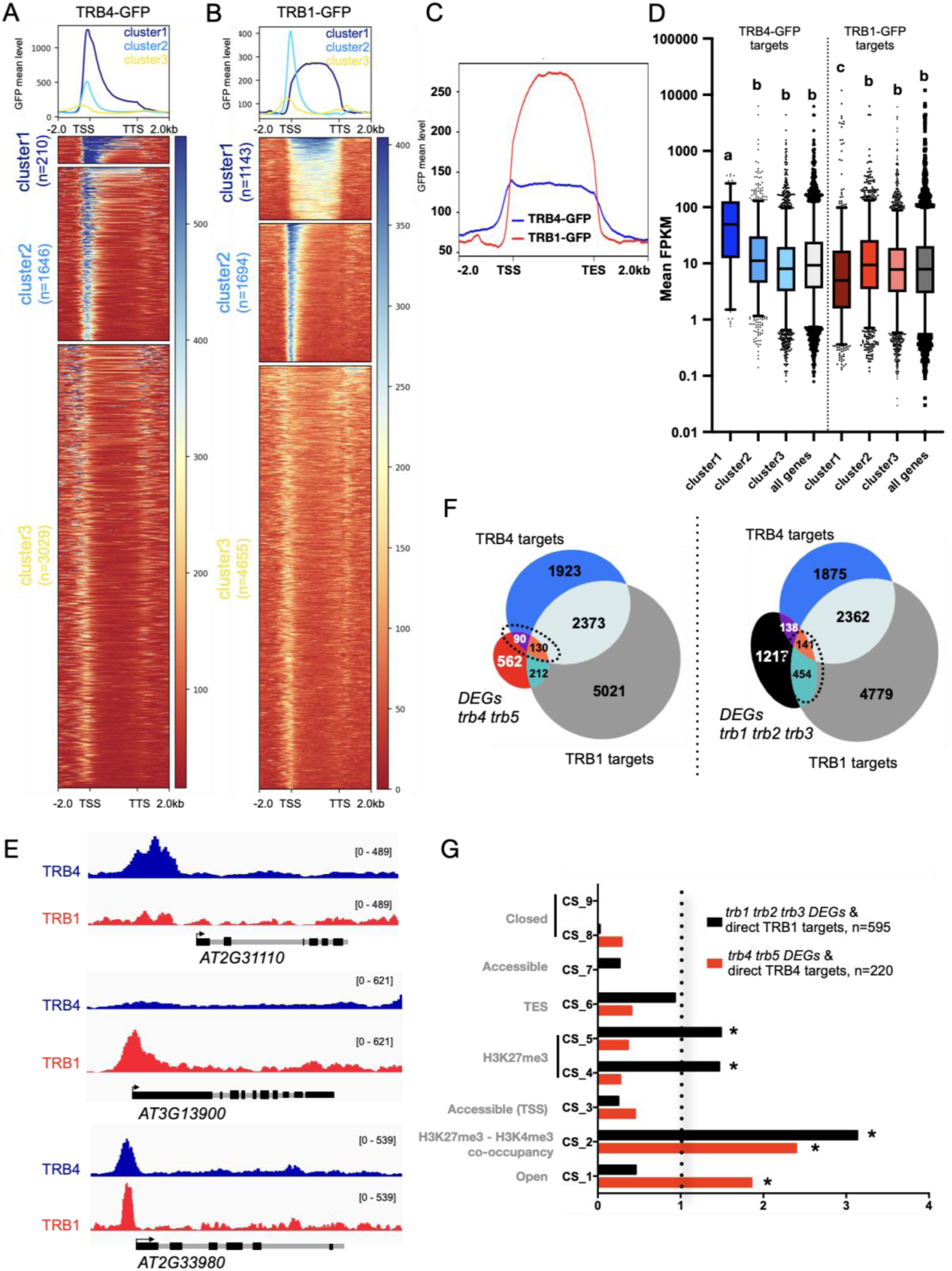
TRB1 and TRB4 are differentially distributed along the genome but share target genes. **(A, B)** Metagene plots and heatmaps after k-mean clustering showing ChIP-seq signals of TRB4-GFP **(A)** or TRB1-GFP **(B)** over TRB4 and TRB1 target genes respectively. **(C)** Metagene plot showing enrichment of TRB1-GFP and TRB4-GFP over TRB1 cluster 1 genes (n=1143). **(D)** Mean expression (FPKM) of genes within the three clusters and for all TRB4-GFP or TRB1-GFP target genes. Different letters indicate significant differences among samples by t-test (p < 0.01). **(E)** Genome browser views of representative genes that are targets solely of TRB4, TRB1 or both. (**F**) Venn diagrams displaying overlap between TRB1 (grey) and TRB4 (blue) targets and DEGs in *trb4 trb5* (left) or in *trb1 trb2 trb3* mutants (right). Overlap TRB4 targets / TRB1 targets (n=2503), p=0; DEGs *trb4 trb5* / TRB4 targets (n=220), p= 3.6e^-09^; DEGs *trb1 trb2 trb3* / TRB1 targets (n=595), p=9.4e^-06^; DEGs *trb4 trb5* / TRB1 targets (n=342), p= 5.6e^-09^; DEGs *trb1 trb2 trb3* / TRB4 targets (n=279), p=0.98. Candidate target genes (220 for TRB4/*trb4 trb5* – 595 for TRB1/*trb1 trb2 trb3*) are delineated **(G)** Enrichment of *trb4 trb5* (n = 220) and *trb1 trb2 trb3* DEGs (n=595) that are direct targets of the respective TRB protein in the nine chromatin states defined by (Sequeira-Mendes et al., 2014) (*OR>1 and p<0.01).

Given the ability of TRB1 and TRB4 proteins to from hetero-dimers ((Kusová et al., 2023), ***Figure 3A-B***) and the faculty of TRB1 and TRB4 to bind similar consensus sequences, we searched for potential TRB1 and TRB4 co-occurrence. In addition to TRB1- and TRB4-specific target sites, TRB4 shares more than half of its binding sites with TRB1 (***Figure 5E -F***). Comparison of the TRB1 and TRB4 targets identified by ChIP-seq with the list of genes misregulated in *trb4 trb5* or *trb1 trb2 trb3* mutant lines identified a few hundred plausibly directly regulated TRB1 or TRB4 targets (***Figure 5F***). Half of the misregulated genes (n=220) that are directly targeted by TRB4 is up-while the other half is down-regulated, suggesting that TRB4 can act either as a positive or as a negative regulator of transcription, potentially depending on the genomic context or on distinct protein interactions. Comparison of *trb4 trb5* mutant DEGs with TRB1 targets also revealed a small, but significant number of genes either targeted commonly by TRB4 and TRB1 or only by TRB1 but not TRB4 (***Figure 5E, F***). These observations suggest the existence of distinct TRB complexes where TRB1 and TRB4 often bind to common genes and potentially influencing each other’s function to regulate gene expression.

### TRB4 is enriched at H3K4me3-marked genes but does not affect H3K4me3 deposition

Closer investigation of the chromatin states associated with the genes misregulated in the *trb4 trb5* or in the *tr1 trb2 trb3* mutant plants and directly bound by either TRB4 or TRB1, revealed that TRB1 targets are overrepresented among genes corresponding to the reference CS2, CS4 and CS5 chromatin states, which are enriched in H3K27me3 and mainly comprise silent or low expressed genes in public datasets ^43^ (***Figure 5G***). Instead, TRB4-associated genes are over-represented among genes carrying both H3K27me3 and H3K4me3 (CS2) or CS1 that usually encompasses active genes with strong H3K4me3 enrichment (***Figure 5G***).

To gain insight in the chromatin marks present at TRB4 and TRB1, we carried out H3K4me3 and H3K27me3 ChIP-seq at the same developmental stage as in our *trb4 trb5* RNA-seq analysis and plotted the distribution of TRB4 and TRB1 as a function of presence and/or absence of these post-translational modifications. Our ChIP-seq profiles show that while TRB1 was expectedly associated with the body of genes marked by H3K27me3 or by both H3K4me3 and H3K27me3, TRB4 is excluded from H3K27me3-marked gene bodies but moderately enriched at TSS and TTS (***Figure 6A***).

**Figure 6:**
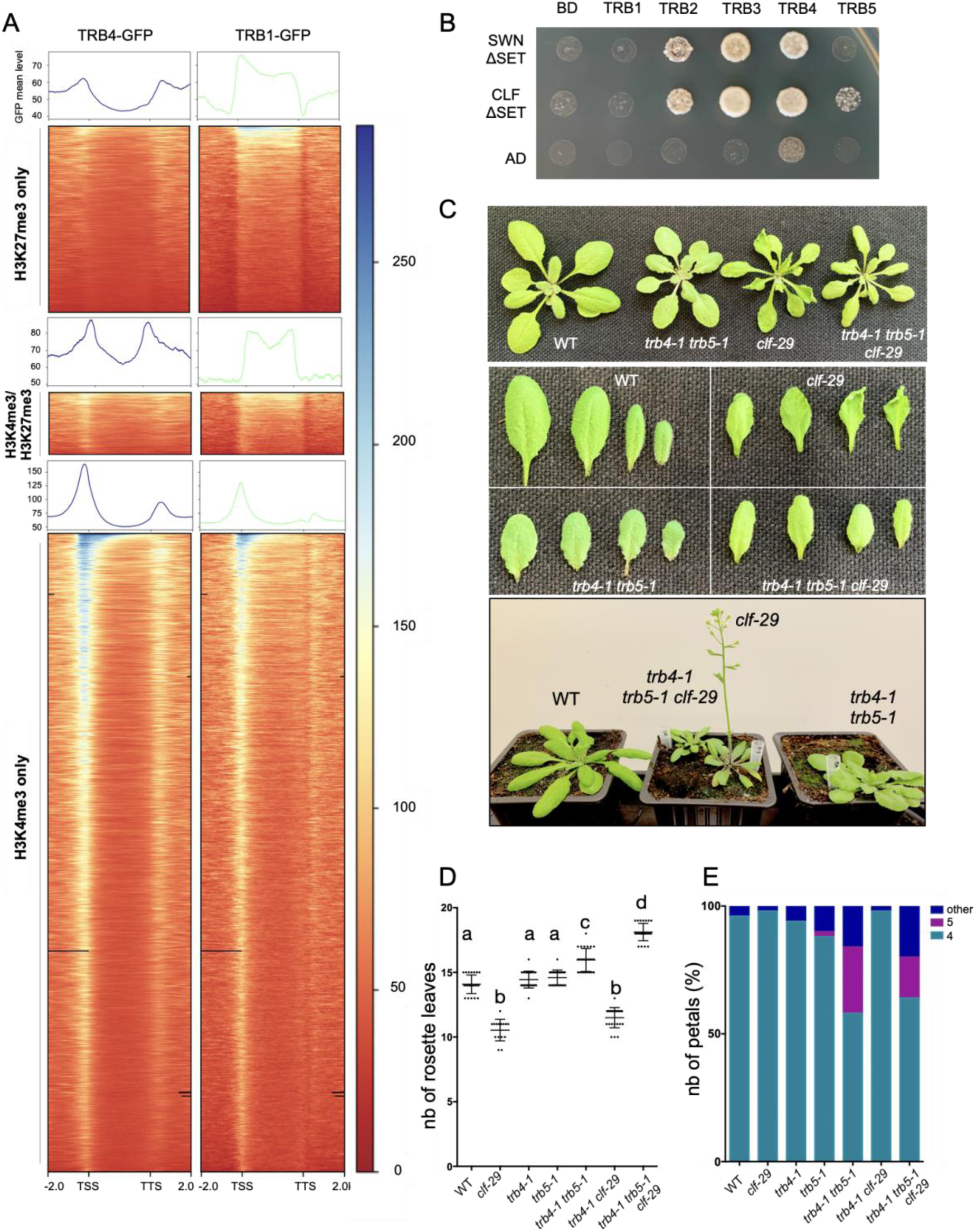
TRB4 *and* TRB5 are required for leaf curling and early flowering in clf mutant plants. **(A)** Metagene plots and heatmaps showing ChIP-seq signals of TRB4-GFP or TRB1-GFP over genes enriched in H3K27me3, H3K4me3 or both histone marks as determined by ChIP-seq analysis. **(B)** Interaction of the five Arabidopsis TRB proteins with CURLY LEAF (CLF) and SWINGER (SWN) proteins lacking the SET domain probed in the Yeast-Two-Hybrid system. Yeast strains growing on synthetic medium lacking Leu, Trp, His and Ade reveal interaction**. (C)** Representative wild type, *trb4-1 trb5-1, clf-29,* and *trb4-1 trb5-1 clf-29* triple mutant plants at 3 weeks (top - middle) and 4 weeks (bottom) after sowing. Loss of TRB4 and TRB5 in the *clf-29* mutant background abolishes the curly leaf and the early flowering phenotype. **(D)** Mean number of rosette leaves at bolting in the indicated genotypes. **(E)** Mean number of petals observed in flowers from the indicated genotypes.

While TRB1-GFP and TRB4-GFP mean profiles at genes differ, with TRB4-GFP being typically enriched at the 5’ and 3’ ends of genes, both proteins mark the TSS of genes associated with H3K4me3 (***Figure 6A, Supplementary Figure 6A***). Considering these observations and the demonstrated role for TRB1-3 proteins in H3K4me3 removal (Wang et al., 2023), we tested whether loss of TRB4/5 affects H3K4me3 enrichment genome-wide. Immunostaining (***Supplementary Figure 6B***) and H3K4me3 ChIP-seq revealed that H3K4 trimethylation patterns are globally unaffected in young *trb4 trb5* plantlets (***Supplementary Figure 6C***). Only a few genes (n=120) show significant changes in this histone mark. Furthermore, plotting H3K4me3 specifically at TRB4 direct targets confirms that the enrichment in this mark is generally maintained in the double mutant (***Supplementary Figure 6D***). Therefore, in contrast to the increased level of H3K4me3 reported in the *trb1 trb2 trb3* mutant (Wang et al., 2023), and despite the enrichment of TRB4 at genes marked by H3K4me3, loss of TRB4 and TRB5 does not affect H3K4me3 levels at most TRB4 binding sites, suggesting that modulating H3K4 methylation or demethylation is not their major mode of action.

### Loss of TRB4 and TRB5 rescues leaf curling and precocious flowering defects in *clf* mutants

TRB1-3 proteins have previously been shown to directly interact with CLF and SWN (Zhou et al., 2018), (Kusová et al., 2023) *via* their coiled-coil domain. Since TRB4 and TRB5 differ in their coiled-coil domain region from clade I TRBs (***Figure 1D, Supplementary Figure 1B, C***), we tested whether they could still be engaged in similar interactions. In agreement with (Kusová et al., 2023), TRB4 interacts with both CLF and SWN in Y2H assays (***Figure 6B***), suggesting that TRB4 could indeed recruit PRC2 to chromatin similar to clade I TRBs or, alternatively, compete with TRB1 for PRC2 interaction. To investigate whether TRB4 interaction with CLF and SWN regulates H3K27me3 enrichment, we profiled the genome-wide distribution of H3K27me3 in 7-day old plantlets. Most genes retain wild-type levels of this histone mark in *trb4 trb5* mutant plants, but around 200 genes (2.7%) show significant gain or loss in H3K27me3 levels (***Supplementary Figure 6E-G***). Hence, compared to the *trb1 trb2 trb3* mutant where 22% of the H3K27me3 enriched genes show altered H3K27methylation (Zhou et al., 2018), the absence of TRB4 and TRB5 affects H3K27me3 at a smaller subset of genes at this developmental stage.

In line with the critical function for TRB1-3 in H3K27me3 deposition, loss of TRB1 and TRB3 induces an aggravation of the single *clf-28* mutant phenotype (Zhou et al., 2018). To investigate the relationship between PcG function and TRB4 and TRB5, we crossed the double mutant with *clf-29* (Schönrock et al., 2006). Surprisingly and in contrast with what was seen with *trb1* mutants (Zhou et al., 2018), several of the *clf-29* phenotypical characteristics were rescued by removal of TRB4 and TRB5. The triple mutant plants showed neither downward curled leaves nor early flowering; instead, flowering was further delayed compared to the *trb4 trb5* double mutant (***Figure 6C, D***). Based on our RNA-seq analysis, this observation is not an indirect consequence of misregulation of the genes encoding the subunits of the PRC2 and PRC1 complexes, as none of the major protein-coding genes of these complexes are misexpressed in the *trb4 trb5* double mutant. (***Supplementary Table 4***). While *clf-29* phenotypic defects were rescued or reverted by *trb4 trb5*, the *CLF* mutation did not reciprocally revert developmental defects specific to the *trb4 trb5* double mutant such as altered leaf color and the supernumerary petal phenotype (***Figure 6E*).** These clade II TRB phenotypes are therefore likely caused by PcG independent processes and tend to be dominant over *clf-29* phenotypes. Therefore, TRB4 and TRB5 physically interact with CLF and SWN two major H3K27 histone methyltransferases, influence H3K27me3 at a small set of genes and are required for the leaf morphology and flowering defects characteristic for the *clf* mutant.

### TRB4 and TRB5 function as transcriptional activators of *FT* and *SOC1*

Flowering time control involves a complex regulatory network that integrates endogenous factors and environmental cues and requires the interplay of chromatin modifications including the PcG pathway and transcription factors at key flowering regulator gene. Therefore, to gain insight into the complex relationship between PRC2-CLF complexes and clade II TRBs in the control of flowering regulators, we investigated the expression of *SEPALLATA3* (*SEP3), FLOWERING LOCUS T (FT)* and *SUPPRESSOR OF OVEREXPRESSION OF CONSTANS 1 (SOC1)* that are targets of TRB4 and *FLOWERING LOCUS C (FLC)* and *AGAMOUS (AG)* not bound by TRB4. From our RNA-seq analysis in young seedlings, *SOC1* is significantly downregulated in the *trb4 trb5* double mutants while *FLC* is upregulated. As *soc1* mutant plants display delayed flowering (Samach et al., 2000), and FLC is a flowering repressor (Michaels and Amasino, 1999; Sequeira-Mendes et al., 2014), *SOC1* down- and *FLC* up-regulation in *trb4 trb5* plants could at least partly explain the late flowering phenotype of *trb4 trb5* mutant (***Figure 2D, Supplementary Figure 2E***). Expression of the other tested genes is either unaffected or too low to be detected at this developmental stage (***Figure 7A***). These observations concord with the tight repression of flowering controlling genes in young tissues, which is illustrated by H3K27me3 enrichment over their gene bodies (***Figure 7A)*.**

**Figure 7:**
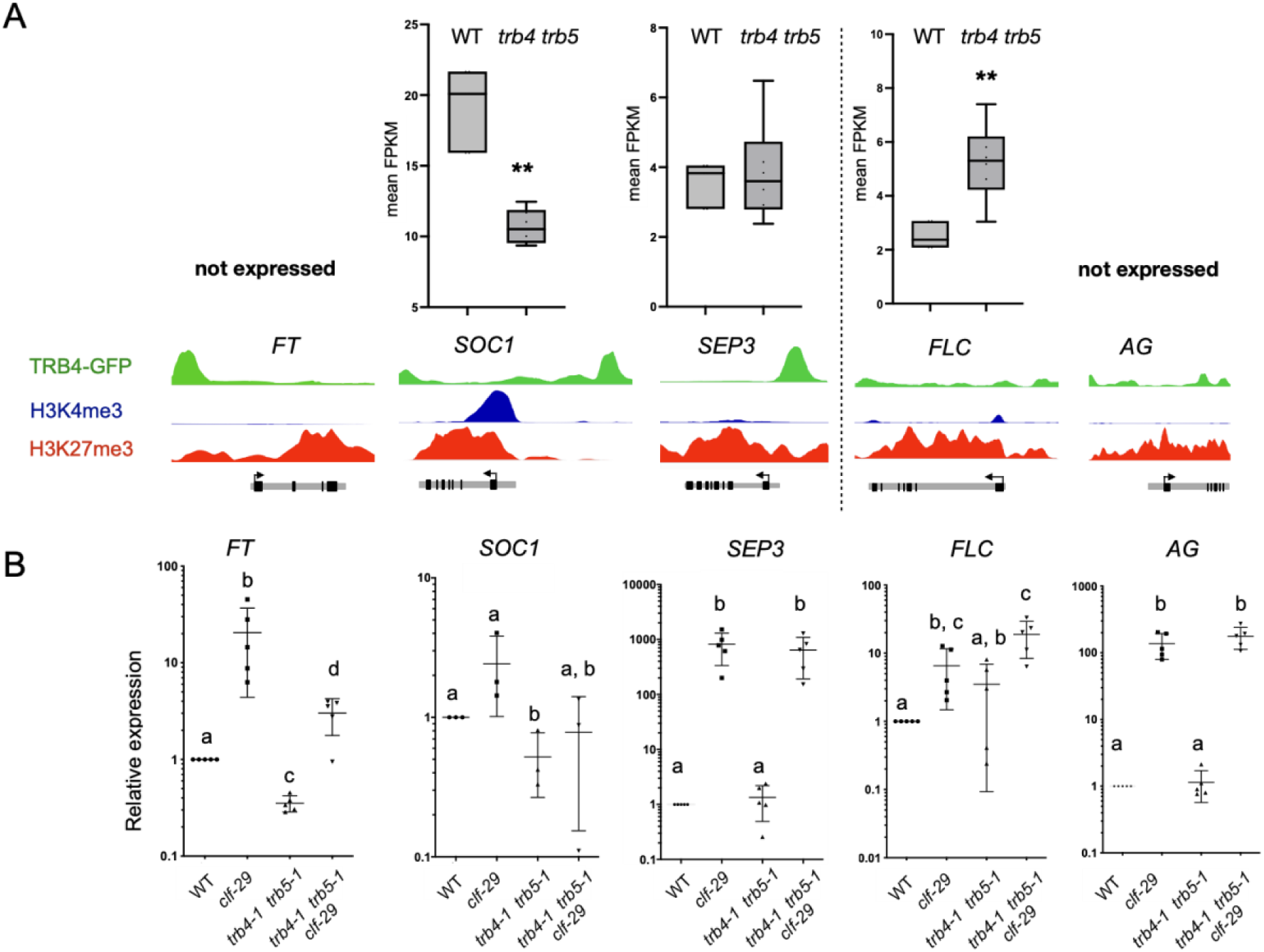
TRB4 and TRB5 function as transcriptional activator of FT and SOC1 flowering regulators. **(A)** *Top:* Expression (FPKM) of *FLOWERING LOCUS T (FT), SUPPRESSOR OF OVEREXPRESSION OF CO 1* (*SOC1*), *SEPALLATA3 (SEP3), FLOWERING LOCUS C (FLC)* and *AGAMOUS (AG)* in RNA-seq datasets from wild type and *trb4 trb5* mutant 7-days old seedlings. Asterisks indicate significant differences among samples by the Mann-Whitney test (p < 0.01). *Bottom:* Genome browser views showing binding of TRB4 and enrichment in H3K4me3 and H3K27me3 at these genes as determined by ChIP-seq at the same developmental stage. *SEP3, FT* and *SOC1* are TRB4 target genes. **(B)** Relative transcript levels determined by RT-qPCR of *FT*, *SOC1, SEP3, FLC* and *AG* in rosette leaves from wild type, *trb4-1 trb5-1, clf-29,* and *trb4-1 trb5-1 clf-29* triple mutant plants at 3 weeks of age. Different letters indicate significant differences among samples by the Mann-Whitney test (p < 0.01).

We then tested how transcript levels of these genes are affected at later developmental stages and in the three mutant conditions (*clf*, *trb4 trb5* and the triple *trb4 trb5 clf* mutant). We therefore extracted RNA from mature leaves of plants before bolting, when these flowering regulators are expressed and determined relative transcript levels by RT-qPCR. In agreement with the tight control by the PcG machinery (Goodrich et al., 1997; Jiang et al., 2008; Lopez-Vernaza et al., 2012) all five genes are upregulated in the *clf-29* mutant. Out of the three direct TRB4 targets, *FT* and *SOC1* transcript levels are reduced in *trb4 trb5* compared to WT (***Figure 7B-C***). Furthermore, in the triple mutant, loss of TRB4 and TRB5 attenuates the transcriptional activation of *FT* and *SOC1* observed in absence of CLF (***Figure 7B-C***), revealing that clade II TRB proteins function as transcriptional activators of *FT* and *SOC1* both in wild type plants and in the *clf-29* background (***Figure 7C***). Since *FT* loss of function is sufficient to revert *clf* early flowering and leaf curling (Lopez-Vernaza et al., 2012), downregulation of this master flowering time regulatory gene in *trb4 trb5 clf* could be causative for the phenotypic suppression of the *clf* flowering time phenotype (***Figure 7C***).

Taken together our observations point to a multifaceted interaction of clade II TRBs with other members of the TRB family and with the PcG machinery mediating developmental and growth control including flowering time regulation. Our results identify TRB4 and TRB5 proteins as novel transcriptional regulators of the floral integrators *FT* and *SOC1* and the flowering repressor *FLC* and reveal their role in fine-tuning flowering time.

## Discussion

### TRB4 and TRB5 do not play a major role in telomere protection

The five Arabidopsis TRBs, including TRB4 and TRB5 comprise two DNA-binding domains: an N-terminal Myb/SANT domain and a central GH1 domain (Kotlinski et al., 2017). In humans, the Myb/SANT domain, which mediates interaction with telomeric double-stranded DNA can be found in only two proteins: TRF1 and TRF2, which form the core of the “shelterin” complex, a complex of six proteins dedicated to telomere protection (Palm and Lange, 2008). In *Arabidopsis thaliana*, in addition to the five TRBs, twelve other proteins harbor the Myb/SANT domain (Schrumpfová et al., 2019) and despite numerous studies, no true “shelterin” has yet been isolated in plants. For example, the absence of TRB1-3 ((Zhou et al., 2018), our unpublished data) or of the six TRF-like proteins (Fulcher and Riha, 2016) does not lead to telomere deprotection. We initially identified TRB4 and TRB5 in a pull-down experiment designed to identify telomeric DNA-binding proteins, and their binding capacity to telomeric repeats was confirmed by EMSA assays (Kusová et al., 2023). This study also revealed that TRB4 and TRB5 can interact with the catalytic subunit of the telomerase (TERT) as well as several telomerase interacting proteins (POT1Aa POT1b, RUVBL1 and RUVBL2A), suggesting that TRB4 and TRB5 are part of the telomerase complex, as has been shown for TRB1-3 proteins (Schrumpfová et al., 2014). However, deletion of TRB4 and TRB5 does not affect telomere protection. Therefore, either deletion of only certain members of the TRB family is not sufficient due to functional redundancy or true shelterin proteins still remain to be identified.

While telomere protection is ensured even in the absence of either clade I or clade II TRBs, plant development and gene expression are affected in the *trb1 trb2 trb3* and in the *trb4 trb5* mutants suggesting a role for TRB proteins beyond telomere function. A role for telomeric proteins beyond chromosome end protection first came to light about 30 years ago with the demonstration of transcriptional silencing of genes located near telomeres in humans and yeast, a mechanism called TPE for Telomere Position Effect or TPE-OLD (Over Long Distance) for silencing of more distant genes (Gottschling et al., 1990; Robin et al., 2014). ZBTB48 one of the most conserved factors associated with human telomere acts as a negative regulator of telomere length but also acts as a transcriptional activator, regulating the expression of a defined set of target genes (Jahn et al., 2017). More recently, in mammals, TRF2 has been shown to bind to short telomeric sequences present in the promoters of certain genes, to participate in the deposition of active marks (H3K4me1 and H3K4me3) as well as silencing marks (H3K27me3), and to affect the transcription of these genes (Simonet et al., 2011; Mukherjee et al., 2018; Mukherjee et al., 2019). While the mechanism behind this control is still unclear and affects only a small proportion of genes, the control of gene transcription by telomeric factors appears to be a conserved function among eukaryotes.

### TRBs origin and evolution

TRB proteins are plant-specific proteins that appeared early in plant evolution as shown by our phylogenetic analyses and those of other’s (Kotlinski et al., 2017; Kusová et al., 2023). In an ancestor of spermatophytes, the TRB protein family split into two clades comprising either Arabidopsis TRB1-3 (clade I) or TRB4-5 (clade II). Although all TRB proteins share a global common architecture with three major domains, we have revealed divergences between the two clades and our results argue for a functional specialization of these clades. The TRB clades are mainly distinguished by their C-terminal coiled-coil domains. This domain, which mediates the interaction of clade I TRBs with the catalytic subunits of the PRC2 complex (Zhou et al., 2018; Kusová et al., 2023), is probably also responsible for the interaction of TRB4 and TRB5 with CLF and SWN, but may allow TRB4 and TRB5 to bind additional, specific partners that remain to be discovered. In almost all angiosperm genomes with the sole exception of *A. comosus,* we found at least one member of each clade, suggesting a requirement for balance between the different functions performed by members of the two TRB clades.

### TRBs interaction with each other and with DNA

Expression analysis does not indicate any tissue-specific expression of the five *TRB* genes during plant development, except for the higher *TRB1* transcript levels observed in embryo and endosperm. Therefore, the different TRB proteins could be present simultaneously in a given cell. Our results as well as the work of others (Kusová et al., 2023) revealed that all TRB proteins can physically interact with each other, both in the yeast system and *in planta*. While a single ‘telobox’ motif is sufficient for TRB proteins to bind *in vitro* (Kusová et al., 2023), TRBs may bind to DNA simultaneously as multimers or compete with each other for the same sites. Given that TRB hetero- and homodimer formation is likely to occur via the GH1 domain as demonstrated for TRB1 homodimerization (Schrumpfová et al., 2014), we can postulate that certain genomic sites are co-bound by TRB proteins from both clades. TRB1 or TRB4 could bind DNA via its Myb/SANT domain and interact with another TRB protein via its GH1 domain. Alternatively, several TRB proteins could bind to the same gene via multiple telobox motifs or via other DNA motifs bound by the GH1 domain.

Finally, binding to the same sites but at different time points or in different tissues, which is not resolved by bulk-tissue ChIP-seq analysis, could also be consistent with the observed co-occurrence of TRB1 and TRB4. Our ChIP-seq experiments show that TRB proteins from the two clades have not only common but also specific targets. Therefore, small differences in their respective DNA-binding domains, the chromatin environment or interaction with specific partners may influence their localization on chromatin.

### TRBs have specific and redundant roles

If clade I and clade II TRB proteins coordinately regulate gene expression, we would expect a set of DEGs common to *trb1 trb2 trb3* and *trb4 trb5* mutants, which is the case for a fraction of them (**Figure 2E**). However, most of the misregulated genes are specific to each mutant suggesting both common and specific roles for each clade.

Previously, it was suggested that all TRB clade I proteins have redundant functions. Indeed, TRB1-3 are highly co-localized throughout the genome (Wang et al., 2023) and the severe developmental defects observed in the triple *trb1 trb2 trb3* mutants can be complemented by any of the TRB clade I proteins ^25^. We show here that while the *trb2 trb3 trb4 trb5* quadruple mutant is viable, the *trb1 trb4 trb5* triple mutant is not, illustrating a specific function of TRB1 that cannot be fulfilled by TRB2 or TRB3 in absence of TRB4 and TRB5. A specific role for TRB1 is supported by the fact that TRB1 is present in all the dicots analyzed, whereas TRB2 and TRB3, which have appeared after a more recent duplication, are absent in several plant species such as tomato (**Figure 1**). Overall, our results highlight that proteins from the two clades may share essential roles, but further analyses are required to decipher whether the observed lethality of the *trb1 trb4 trb5* triple mutant is related to a defect in gene transcriptional control at a critical embryonic stage or to some other reason. A scenario is therefore emerging, in which proteins from the two different clades work together or play opposing roles in coordinating the expression of target genes. Identifying both the physical and genetic interactors of each TRB may shed light on their specific functions.

### Clade II TRB proteins function as transcriptional regulators

Clade I TRB proteins have been described to recruit histone modifiers (PRC2, JMJ14) to silence a subset of developmental genes by participating in H3K27me3 deposition and H3K4me3 removal (Zhou et al., 2018; Wang et al., 2023) at promoters containing telobox DNA motifs. Consistent with this function, loss of all three clade I TRB proteins results in developmental growth defects similar to severe PRC2 mutants and loss of one clade I TRB protein alone is sufficient to enhance *clf* mutant phenotypes ^23^.

Despite their ability to interact with CLF and SWN (**Figure 3**) as well as EMF2 and VRN2 (Kusová et al., 2023), double mutant plants lacking both clade II TRBs do not show such PRC2 mutant like phenotype, but harbor milder developmental phenotypes such as late flowering or supernumerary petal numbers that have been reported in mutants deficient in H3K27me3 removal (Carles and Fletcher, 2009; Yan et al., 2018) or H3K4me3 deposition (Alvarez-Venegas et al., 2003) suggesting that TRB4 and TRB5 counteract PRC2 silencing activity at certain genes. In agreement with this hypothesis, some phenotypes associated with PRC2 deficiency (early flowering, curly leaves) are restored to normal in the triple *trb4 trb5 clf-29* mutant plants. Based on these genetic data that argue for a role of clade II TRBs in counteracting PRC2 activity, we expected to observe altered H3K27me3 and/or H3K4me3 homeostasis but these marks were only affected at a small subset of genes upon loss of TRB4 and TRB5 and only few showed changes in gene expression. Altogether this suggests that modulating the H3K27me3/H3K4me3 balance is not the major mode of action of clade II TRB proteins.

Instead, the phenotypic restoration of the *clf* mutant phenotype could be indirect, by altering the levels of other factors involved in chromatin regulation, or direct by modulating the expression of specific genes that are targets of CLF such as FT. Loss of *FT* an essential developmental integrator, in the *clf* background has previously been shown to be sufficient to suppress early flowering and leaf curling, without affecting *AG* expression (Lopez-Vernaza et al., 2012). Indeed, in mature leaves, transcript levels of *FT* and *SOC1,* which are H3K27me3 marked and direct targets of both CLF (Lopez-Vernaza et al., 2012) and TRB4 (this study), are increased in the *clf-29* mutant background, but closer to wild-type levels in the triple mutant. Therefore, clade II TRBs emerge as novel transcriptional activators of specific flowering regulators required for fine-tuning flowering time.

A potential mode of action for clade II TRB proteins that could be tested in the future, is the recruitment of histone acetylation or deacetylation activity that holds a prominent place in the transcriptional control of PRC2 target genes and participates in correct gene expression all along plant development (Wang et al., 2014). For instance, CBP, a Histone AcetylTransferases (HAT) that acetylates H3K27 (H3K27Ac) antagonizes Polycomb silencing (Tie et al., 2014). Specifically, plants deficient for HAC1 and HAC5, two HATs from the MEDIATOR complex harbor developmental defects resembling *trb4 trb5* defects: plants are small and show delayed flowering and reduced fertility (Guo et al., 2021). As was demonstrated for clade I TRBs, TRB4 and TRB5 interact also with members of the PEAT complex (Kusová et al., 2023) that is involved in histone deacetylation to silence heterochromatin (Tan et al., 2018) and it has also been reported that TRB2 interacts with HDT4 and HDA6, two histone deacetylases presumably acting in H3K27 deacetylation (Lee and Cho, 2016). Taking these observations together TRB proteins may, beyond their ability to influence the deposition/removal of H3K27me3 and H3K4me3, be involved in the coordination of histone acetylation/deacetylation in a yet to be defined manner.

### The GH1 protein family forms a complex network of proteins

The mode of action of TRB proteins is complexified by the presence of several other proteins also harboring a GH1 domain in *Arabidopsis thaliana*, namely H1 linker histones and GH1-HMGA proteins (Kotlinski et al., 2017). GH1-HMGA proteins that bind 5′ and 3′ ends of gene bodies similar to a subset of TRB4 targets (**Figure 5**) have been implicated in the repression of *FLC* by inhibiting gene loop formation, which facilitates its transcriptional activation (Zhao et al., 2021). Intriguingly, *FLC* is also upregulated in *trb4 trb5* mutants (**Figure 7**). According to their role in transcriptional regulation, TRB proteins may therefore also participate in the regulation of gene loops by forming homo/heterodimers between proteins linked to nearby motifs. Moreover, rice single Myb transcription factor TRBF2 as well as Arabidopsis TRB1 form phase-separated droplets, which aggregate with PRC2 (Xuan et al., 2022) revealing a propensity of TRB proteins to form phase-separated condensates undoubtedly an essential property for their binding to chromatin and their correct activity.

The globular GH1 domain shared between GH1-HMGA1 and H1 proteins can mediate DNA interaction (Bednar et al., 2017) as well as protein-protein interactions (Schrumpfová et al., 2008) leading to a complex network of interaction/competition between all these proteins. Our recent work already point to a competition between TRB1 and H1 proteins (Teano et al., 2022) and between HMGA1 and H1 (Charbonnel et al., 2018) and future studies will certainly elucidate this interaction network and determine its role in gene expression.

## Materials and Methods

### Gene and protein sequences

Orthologs of *Arabidopsis thaliana* (Ath) TRB proteins were collected from the following plant species that best represent the evolutionary history of the green lineage: *Arabidopsis lyrata* (Aly), *Eutrema salsugineum* (Esa), *Schrenkiella parvula* (Spa), *Brassica rapa* (Bra), *Boechera stricta* (Bst), *Capsella grandiflora* (Cgr), *Descurainia sophioides* (Dso), *Diptychocarpus strictus* (Dst), *Euclidium syriacum* (Esy), *Malcolmia maritima* (Mma), *Myagrum perfoliatum* (Mpe), *Rorippa islandica* (Ris), *Stanleya pinnata* (Spi), *Thlaspi arvense* (Tar), *Prunus persica* (Ppe), *Glycine max* (Gma), *Theobroma cacao* (Tca), *Vitis vinifera* (Vvi), *Populus trichocarpa* (Ptr), *Solanum lycopersicum* (Sly), *Oryza sativa* (Osa), *Zea mays* (Zma), *Sorghum bicolor* (Sbi), *Ananas comosus* (Aco), *Musa acuminata* (Mac), *Amborella trichopoda* (Atr), *Nymphaea colorata* (Nco), *Pseudotsuga menziesii* (Pme), *Picea sitchensis* (Psi), *Pinus lambertiana* (Pla), *Gnetum momentum* (Gma), *Marchantia polymorpha* (Mpo), *Physcomitrella patens* (Ppa), *Oestrococcus lucimarinus* (Olu) and *Chlamydomonas reinhardtii* (Cre). Orthologous sequences were obtained using several sources including mmseqs (Hauser et al., 2016), NCBI (tblastn and blastp) and Phytozome 13 (Goodstein et al., 2012) Gymnosperm orthologs were identified from conGenIE.org (Conifer Genome Integrative Explorer (Sundell et al., 2015). Protein accession numbers and sequences used in this study are listed in **Supplementary Table S1**.

### Phylogenetic studies

Phylogenetic trees were constructed using protein sequences either from Brassicaceae only or species representing the whole plant lineage using MAFFT 7.407 (Katoh and Standley, 2013) for multiple alignment IQ-TREE v2.2.0.3 (Minh et al., 2020) with the LG substitution model for tree with 1000 bootstrap replicates. Trees were refined using the Interactive Tree Of Life (ITOL) (Letunic and Bork, 2016). The MEME 5.1.1 (Multiple Em for Motif Elicitation) suite was used for *de novo* motif predictions of TRB protein sequences from Brassicaceae (Bailey, 2021). From a list of orthologous proteins, MEME was parameterized to define 10 motifs, each with a maximum length of 150 amino acids, with the zoops option.

### Plant Material

Single mutant lines *trb1-1* (SALK_025147) *trb2-1* (FLAG_242F11), *trb3-1* (SALK_134641), *clf-29* (SALK_021003) mutants were provided by the Nottingham Arabidopsis Stock Center. The *trb1-1 trb2-1 trb3-1* triple mutant was obtained by crossing. CRISPR/Cas9 technology as in (Fauser et al., 2012) was applied to generate the *trb4* and *trb5* single mutants with a single RNA guide (**Supplementary Table S3**) targeting the first exon of *TRB4 (At1g17520)* and the second exon of *TRB5 (At1g72740*, ***Supplementary Figure 2a***). Two single mutants for each gene with nucleotide insertions were retained that caused premature stop codons (***Supplementary Figure 2a***) and two different double mutants, termed *trb4-1 trb5-1* and *trb4-2 trb5-2,* were generated by crossing.

For construction of *trb* multi-mutants *trb4-1 trb5-1* was crossed with *trb1-1, trb2-1* and *trb3-1* single mutants*. trb4-1 trb5-1 trb2-1* and *trb4-1 trb5-1 trb3-1/TRB3* were then further crossed to obtain *trb4-1 trb5-1 trb2-1 trb3-1* quadruple mutants.

For plant culture in soil, seeds were stratified for 2 days at 4°C in the dark, and plants grown under long day conditions (16h light, 8h dark, 23°C). For RNA- and ChIP-seq experiments, seeds were sterilized in 70% EtOH / 0.01% SDS and seedlings were grown *in vitro* on 1x MS plates containing 1% sucrose. Transgenic plants were obtained by the floral dip method using the *Agrobacterium tumefaciens* strain GV3101 and transgenic progeny selected by Basta or Hygromycin.

### Plant developmental phenotype description

Root length was measured at 3, 5 and 7 days after germination on 3 independent replicates with 100 plants each. Seed number per silique was counted on 15 siliques taken on the principal stem of 5 individual plants. Flowering time was determined by numbering total rosette leaves at bolting (30 plants from 2 independent experiments). The number of petals was counted on 100 flowers from 5 individual plants. t-test was applied to test for significant differences, except for the root length, for which a two-way ANOVA test was applied.

### Telomere Restriction Fragment (TRF) analysis

TRF analysis of telomere length in HinfI-digested genomic DNA was conducted as described previously (Charbonnel et al., 2018).

### Constructs and cloning

All cloning procedures relied on Gateway technology®. For *in planta* complementation of the *trb4-1 trb5-1* and *trb4-2 trb5-2* double mutants, genomic constructs were obtained by PCR amplification from genomic DNA. Constructs containing either the respective endogenous promoter or the *HMGA2* (*At1g48620*) promoter (used for IF and ChIP-seq) were generated and cloned in pDONR vectors.

For Yeast-Two-Hybrid (Y2H) and Bimolecular Fluorescence Complementation (BiFC) constructs, the cDNA of *TRB5* was obtained by RT-PCR. The cDNA of *TRB4* was synthesized (Integrated DNA Technologies, https://eu.idtdna.com/). After initial cloning into pDONR, constructs were recombined into the appropriate expression vectors for Y2H assays (bait vector pDEST-GBKT7 or prey vector pDEST-GADT7), for *in planta* expression (pB7FWG) or for BiFC (pBiFCt-2in1-NN). The list of all plasmids and oligonucleotides used in this study can be found in **Supplementary Table S3**.

### Yeast Two-Hybrid Assay

Yeast cultures were grown at 30°C on YPD or on selective SD media. Bait (pDEST-GBKT7) or prey (pDEST-GADT7) vectors were transformed into *Saccharomyces cerevisiae* strains AH109 Gold and Y187 (Clontech, MATCHMAKER GAL4 Two-Hybrid System) respectively using a classical heat shock protocol (Gietz and Woods, 2002) and grown on selective medium lacking Trp or Leu. The two yeast strains were mated on YPD and diploids selected on SD-Leu-Trp. Protein-protein interactions were detected by growth on high stringency selective medium lacking Leu, Trp, His and Ade. Empty pDEST-GBKT7 or pDEST-GADT7 vectors were used as negative controls.

### Bimolecular Fluorescence complementation (BiFC)

BiFC vectors were transformed into *Agrobacterium tumefaciens* strain GV3101 and Agrobacterium infiltrated into young *Nicotiana benthamiana* leaves as described (Grefen and Blatt, 2012) together with the p19 suppressor of gene silencing to enhance expression (Norkunas et al., 2018).

### Slide preparation and Immunofluorescence staining

For immunostaining of H3K4me3, H3K27me3, γ-H2A.X and detection of TRB-GFP fusion proteins, nuclei from 7-days old seedlings were isolated as described in (Pavlova et al., 2010). Slides were incubated overnight at 4°C with 50μL of primary antibody in fresh blocking buffer (3% BSA, 0.05% Tween 20 in 1 x PBS), washed 3 x 5 min in 1 x PBS solution, and then incubated 2 to 3 h at room temperature in 50 μL blocking buffer containing secondary antibodies. Finally, slides were washed 3 x 5 min in 1 x PBS and mounted in Vectashield mounting medium with 1.5 μg/mL DAPI (Vector Laboratories). Antibodies and dilutions used in this study are reported **in Supplementary Table 3**. For γ-H2A.X immunostaining, root tips from 7-days old plantlets were treated as described in (Amiard et al., 2011) and the foci were counted for 100 nuclei coming from five individual plants of each genotype. For quantification of anaphase bridges, whole inflorescences were treated as described in (Amiard et al., 2011). At least 100 mitoses were counted from five individual plants.

### Image acquisition and analysis

For the BiFC analysis, fluorescence images of transiently transfected *Nicotiana benthamiana* leaves were obtained using an inverted confocal laser-scanning microscope (LSM800; Carl Zeiss). The 488-nm line of a 40-mWAr/Kr laser, and the 544-nm line of a 1-mW He/Ne laser were used to excite GFP/YFP, and RFP (transfection control), respectively. Images were acquired with 20x or 40x objectives. Images of Arabidopsis roots expressing TRB4- or TRB5-GFP or immunostained isolated nuclei were acquired with a Zeiss epifluorescence microscope equipped with an Apotome device using a 20x objective or a 63x oil immersion objective respectively.

### RNA extraction, RT-qPCR and sequencing

7-day old *in vitro* grown plantlets or adult leaves of soil grown 3-4 week-old plants were ground in 2 mL tubes using a Tissue Lyser (Qiagen) twice for 30 sec at 30 Hz before RNA extraction using the RNeasy Plant Mini kit (Qiagen). For RT-qPCR, RNA was primed with oligo(dT)15 using M-MLV reverse transcriptase (Promega, https://france.promega.com). Relative transcript levels were determined with the LightCycler 480 SYBR Green I Master kit (Roche, https://lifescience.roche.com) on the Roche LightCycler 480 after normalization to *MON1* (*At2g28390*) transcript levels, using the comparative threshold cycle method. Primers used for RT-qPCR can be found in ***Supplementary Table 3***.

For RNA-seq analysis, mRNA was sequenced using the DNBseq platform at the Beijing Genomics Institute (BGI Group) to obtain around 20 million 150 bp paired-end, strand specific reads. Differential expression was determined using an *in house* developed pipeline (https://github.com/vindarbot/RNA_Seq_Pipeline). In brief, reads were trimmed using Bbduk (Bushnell, 2014) to remove adapters and low-quality reads. Clean reads were then aligned to the TAIR10 genome, using STAR (v2.7.1). Read counts per gene were generated using featureCounts (v1.6.3) and the differential expression analysis was performed with DESeq2 (Love et al., 2014) with the threshold log2FC>0.5, padj <0,01. GO-term enrichment of differentially regulated genes was carried out with ClusterProfiler (Wu et al., 2021) using all expressed genes in the dataset as background list.

### ChIP and ChIP-seq analysis

ChIP-seq analysis was essentially carried out as described in (Teano et al., 2022). In brief, about 1g of 7-day old *in vitro* grown plantlets were fixed in 1% formaldehyde under vacuum twice for 7 min and then quenched in 0.125 M glycine. Nuclei were isolated and lysed, and chromatin sonicated using the Diagenode Bioruptor (set to high intensity, 3 times 7 cyles (30sec ON / 30 sec OFF) or the S220 Focused-ultrasonicator (Covaris) for 20 min at peak power 110 W, duty factor 5%, 200 cycles per burst for TRB4-GFP and for 5 min at peak power 175 W, duty factor 20%, 200 cycles per burst for histone modifications for obtain mono-nucleosomal fragments. The following antibodies were used for immunoprecipitation: anti-GFP, Invitrogen, #A-111222, anti-H3K27me3, Diagenode, #C15410069, Batch A1818P, anti-H3K4me3, Millipore #04-745. Immunoprecipitated DNA was recovered by Phenol-Chloroform extraction or Zymo ChIP DNA purifications columns and quantified using a Qubit device. Library preparation using the Illumina TruSeq ChIP kit and sequencing was carried out (DNBSEQ-G400, 1×50bp) at the BGI platform. Each ChIP-seq was carried out in two biological replicates.

### Bioinformatics for ChIP-seq analysis

For the TRB4-GFP ChIP, raw reads were pre-processed with Trimagalore to remove Illumina sequencing adapters. Trimmed reads were mapped against the TAIR10 *Arabidopsis thaliana* genome with Bowtie2 using “--very-sensitive” setting. Peaks were called using MACS2 (Zhang et al., 2008) with the command “macs2 callpeak -f BAM -g 1e8 --nomodel --broad –qvalue 0.01 --extsize 100”. Only peaks found in both biological replicates were retained for further analyses (bedtools v2.29.2 intersect). Annotation of genes and TEs was done using HOMER (annotatePeaks.pl). Metagene plots were generated with Deeptools using computeMatrix and plotProfile commands. TRB4-GFP and TRB1-GFP clusters were identified using Deeptools plotHeatmap using the --kmeans setting. Motifs enrichment under TRB4-GFP peaks was performed using STREME version 5.5.0 (Bailey, 2021). The following options were used “--verbosity 1 --oc . --dna --totallength 4000000 --time 14400 --minw 8 --maxw 15 --nmotifs 10 – align center”.

For H3K27me3 and H3K4me3 enrichment analysis, raw reads were aligned with Bowtie2. Peaks of H3K27me3 read density were called using MACS2 (Zhang et al., 2008) with the command “macs2 callpeak -f BAM --nolambda -q 0.01 -g --broad”. Only peaks found in both biological replicates and overlapping for at least 10 % were retained for further analyses. We scored the number of H3K27me3 or H3K4me3 reads overlapping with marked genes using bedtools v2.29.2 multicov and analyzed them with the DESeq2 package (Love et al., 2014) in the R statistical environment v4.1.2 to identify the genes enriched or depleted in H3K27me3 or H4K4me3 in mutant plants (p-value <0.01).

## Data Availability

The genome-wide sequencing generated for this study have been deposited on NCBI’s Gene Expression Omnibus (GEO) with the accession number of GSE236267 (RNAseq *trb4 trb5*, *trb1 trb2 trb3*), GSE237158 (ChIPseq TRB4-GFP), and GSE237185 (ChIPseq H3K27me3 and H3K4me3 in WT and *trb4 trb5*). All other data supporting the conclusions of the study will be available from the corresponding author upon request.

## Author contributions

SA and AVP designed the research, interpreted the data and wrote the manuscript; SA performed most of the experiments with help from LF, LS, SLG, LL, LW, CB, CT and AVP. FB performed IP–MS and interpreted the data. CB, FB, LS and CT performed Bioinformatic data analyses; and reviewed the manuscript.

## Acknowledgements

We thank Yoan Renaud for help and advice with the bioinformatics analysis, Aurore Pardon and Vincent Darbot for technical help. We thank Claire Jourdain and Daniel Schubert for sharing CLF and SWN constructs for Y2H, and Charles White and Maria Gallego for support and advice during the initial stages of this project. We also thank the “CLIC”, the “Anipath histopathologie” and the “plant” platforms of the iGRED.

## Supplementary data

***Supplementary Figure 1:*** *Supplemental information related to Figure 1*

***Supplementary Figure 2:*** *Supplemental information related to Figure 2*

***Supplementary Figure 3:*** *Supplemental information related to Figure 3*

***Supplementary Figure 4:*** *Supplemental information related to Figure 4*

***Supplementary Figure 5:*** *Supplemental information related to Figure 5*

***Supplementary Figure 6:*** *Supplemental information related to Figure 6*

***Supplementary Table 1:***

1A: Protein sequences of species used in **Figure 1B** and Supplementary Figure 1A
1B: Protein sequences of species used in **Figure 1C**

***Supplementary Table 2:***

2A: DEG in trb4 trb5 and trb1 trb2 trb3 mutants
2B: List of genes targeted by TRB4_GFP or TRB1_GFP
2C: List of H3K27me3 and H3K4me3 genes in WT and in trb4 trb5 mutants

**Supplementary Table 3:**

3A: List of oligos used in this study
3B: List of vectors used in this study
3C: List of antibodies used in this study

**Supplementary Table 4:** FPKM of PRC1 and PRC2 genes in WT and *trb4 trb5* mutants

**Supplementary Figure 1:**
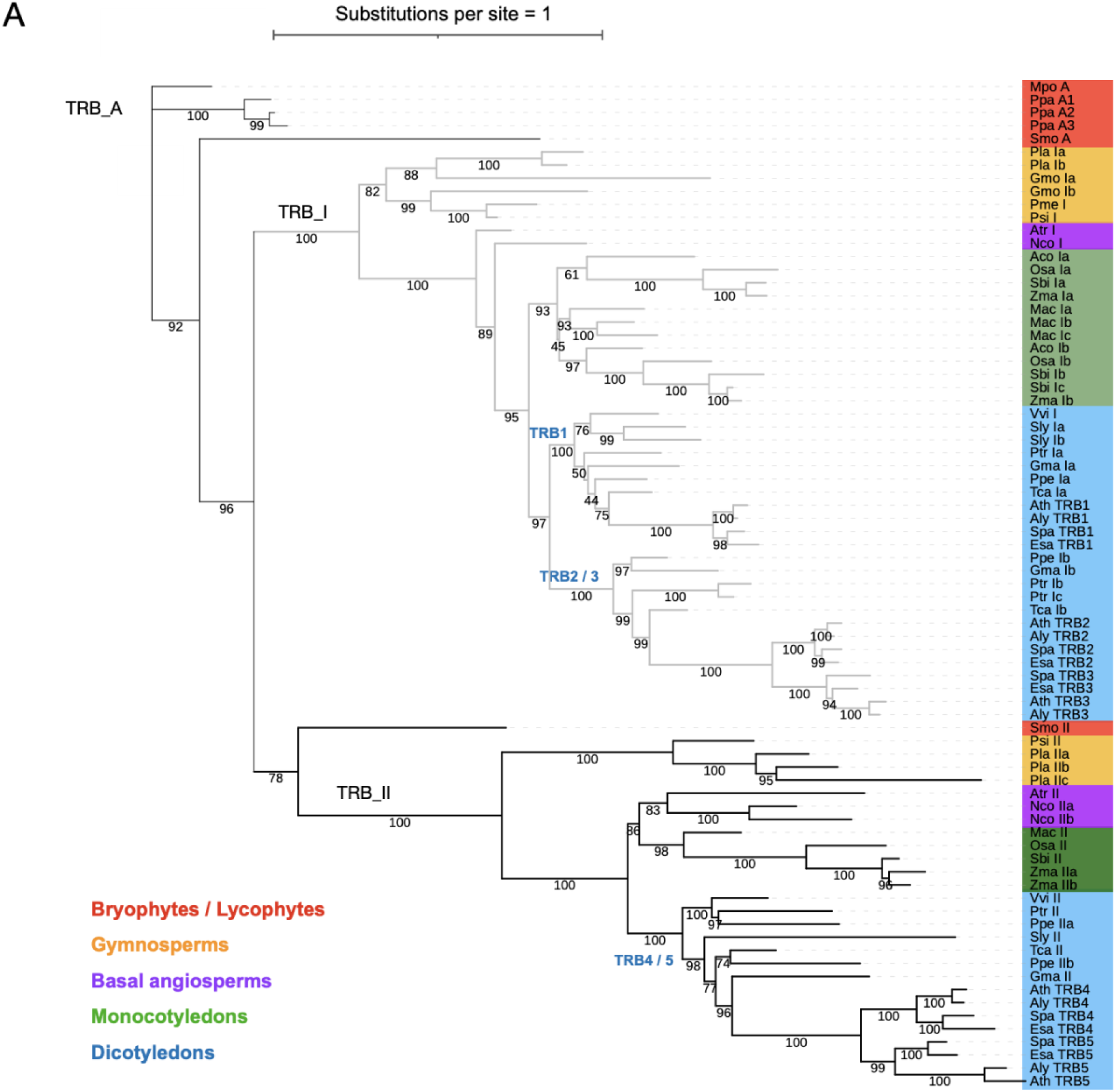
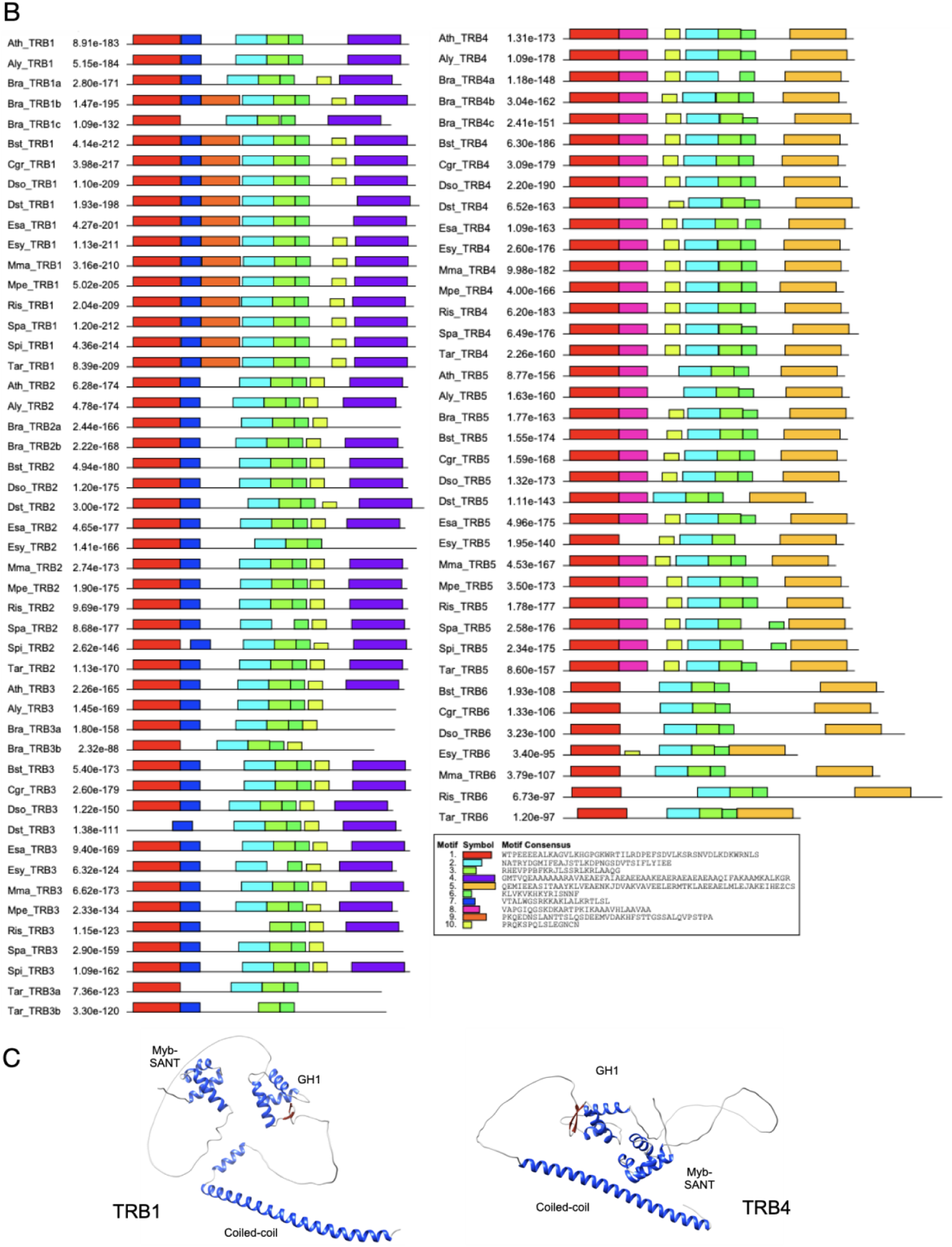
Supplemental information related to Figure 1. **(A)** Rooted maximum likelihood phylogenetic tree for TRB orthologs from 24 plant species. Bootstrap values are indicated for each branch. An ancient TRB clade (TRB_A), a clade comprising Arabidopsis TRB1, TRB2 and TRB3 (TRB_I) and a third clade comprising TRB4 and TRB5 (TRB_II) were defined. **(B)** MEME protein motif prediction of the 10 best motifs among the 15 Brassicaceae species. **(C)** Alpha-fold predictions of long alpha helices in the C-terminal domains of Arabidopsis TRB1 and TRB4 proteins.

**Supplementary Figure 2:**
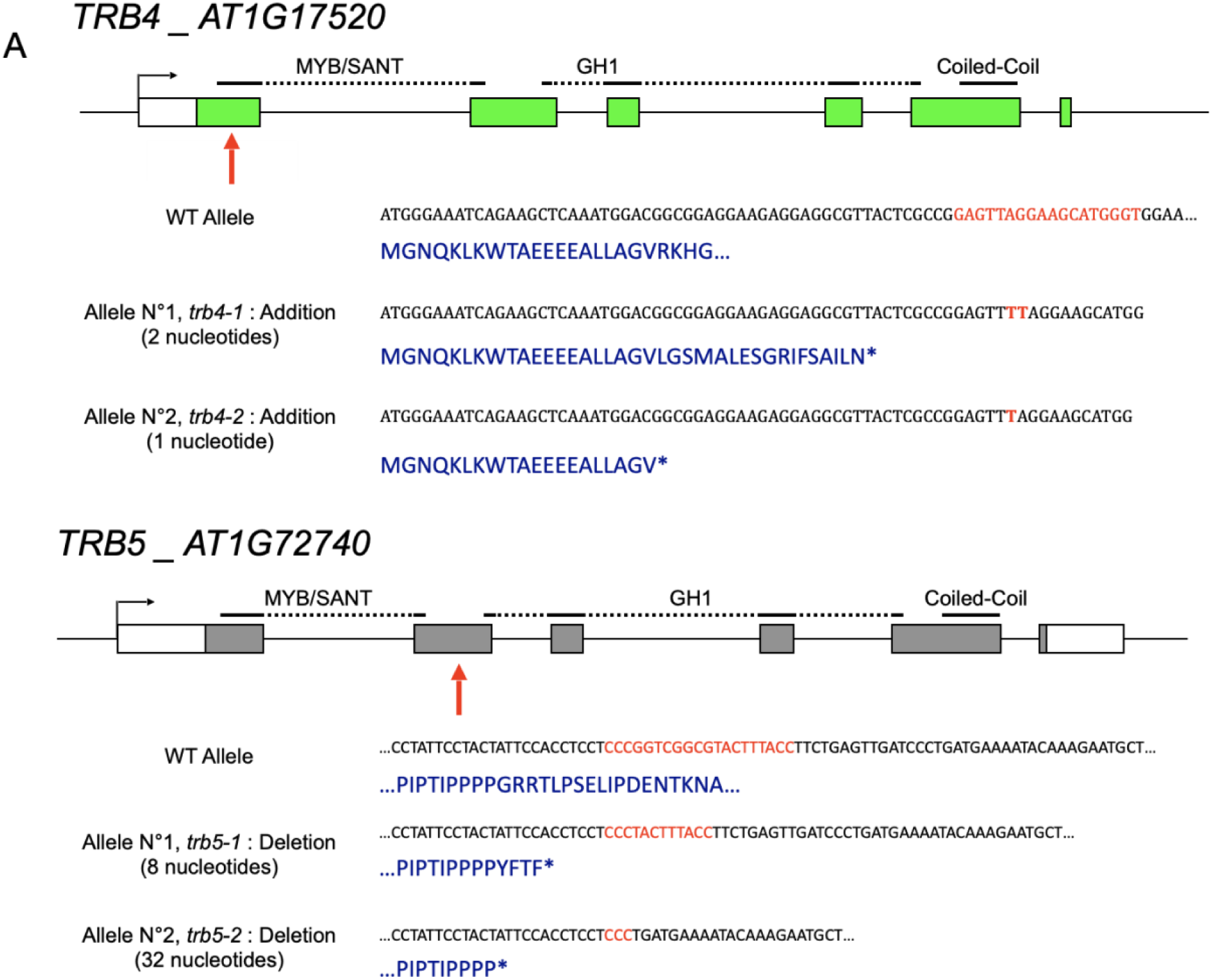
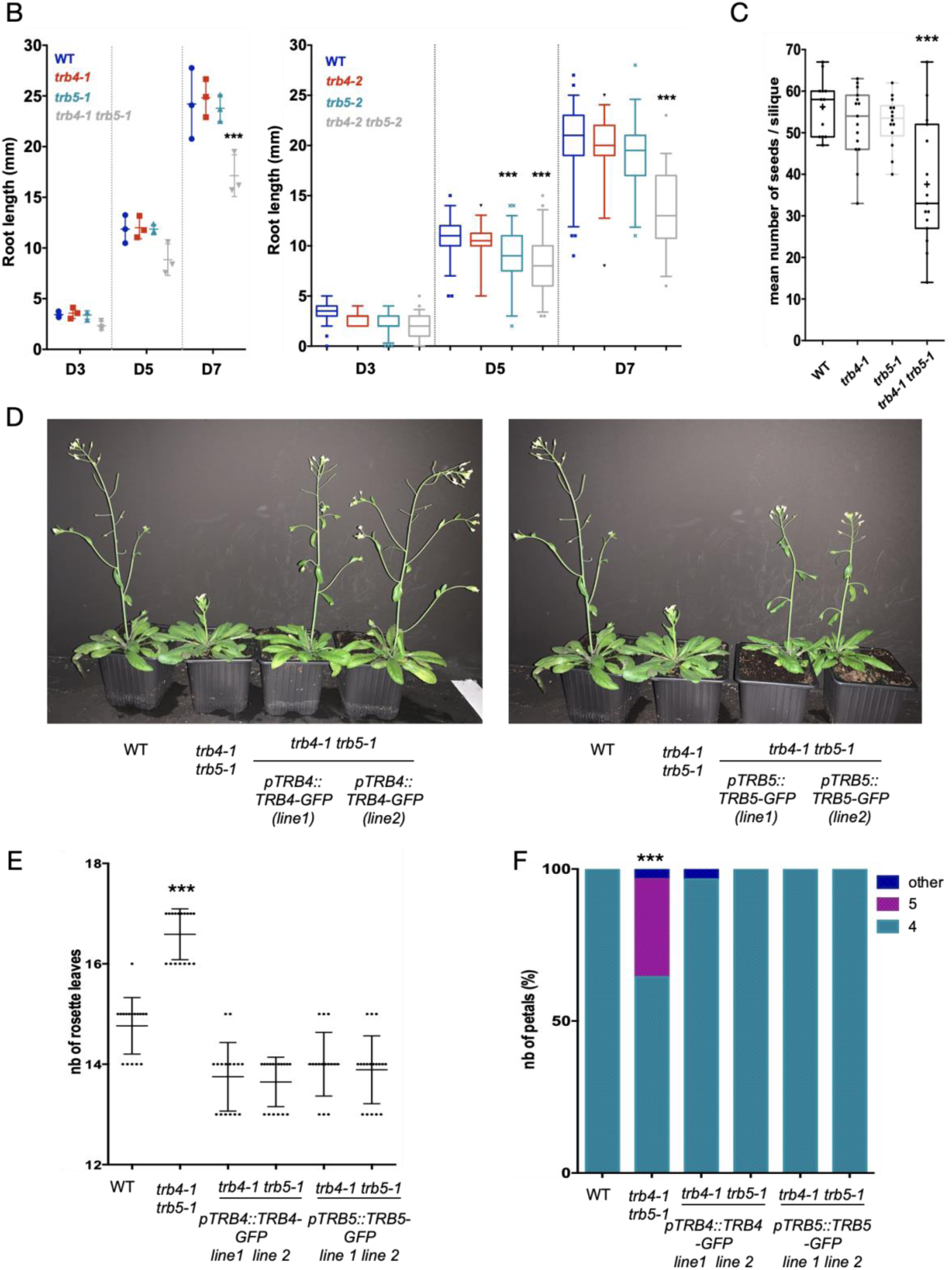
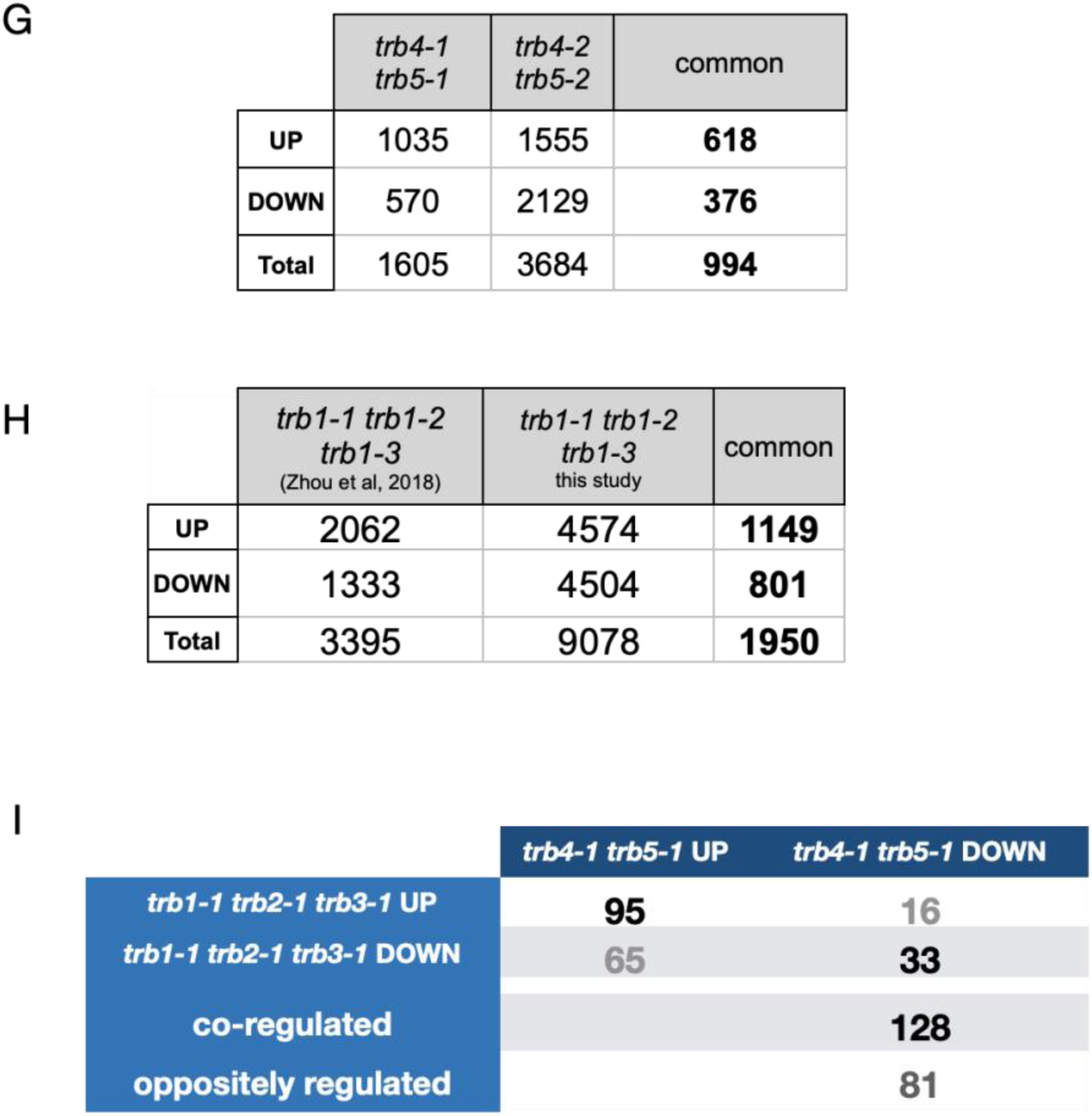
Supplemental information related to Figure 2. **(A)** *TRB4* and *TRB5* mutant alleles generated by CRISPR/Cas9. The corresponding nucleotide and the resulting amino acid sequences are shown. In all mutants, premature stop codons are induced. The red arrow indicates the Cas9 target site. **(B)** Quantification of root length from *in vitro* grown wild type, *trb4-1* and *trb5-1* single and *trb4-1 trb5-1* double mutant plantlets (left) and for wild type, *trb4-2* and *trb5-2* single and *trb4-2 trb5-2* double mutants (right) at day 3 (D3), 5 and 7. For each time point, means from three (left) or two (right) replicates comprising 100 plants each are shown. Roots of *trb4-1 trb5-1 or trb4-2 trb5-2* double mutants are significantly shorter (***p < 0.0001, t-test) at D7. **(C)** Mean number of seeds from 15 siliques of five wild type plants, *trb4-1* and *trb5-1* single and *trb4-1 trb5-1* double mutants. Double mutants are significantly less fertile (***p = 0.0004, t-test). **(D)** Representative 4-weeks-old wild type, *trb4-1 trb5-1* double mutants and four independent transgenic lines expressing either TRB4-GFP or TRB5-GFP under their respective endogenous promoter. The delayed flowering phenotype of *trb4-1 trb5-1* double mutants is complemented. (**E**) Quantification of leaf number at bolting in wild type, *trb4-1 trb5-1* double mutants and four independent transgenic lines shown in (**D**). n = 17, N=1, *** < 0.001, t-test. (**F**) Percentage of flowers showing 4, 5 or any other aberrant petal number in the same genotypes as in (**D**). n=100, N = 5, *** < 0.001, t-test. (**G**) Number of up- and down regulated genes relative to WT in RNA-seq analysis from 3 replicates of *trb4-1 trb5-1* and *trb4-2 trb5-2* mutants. FC > 0.5, padj < 0.01. **(H)** Number of up- and down regulated genes relative to WT in RNA-seq analysis from 3 replicates of *trb1-1 trb2-1 trb3-1* in our data set and the dataset from (Zhou et al., 2018). FC > 0.5, padj < 0.01. **(I)** Comparison of co- or oppositely regulated genes in *trb4 trb5* and *trb1 trb2 trb3* datasets from (**G**) and (**H**).

**Supplementary Figure 3:**
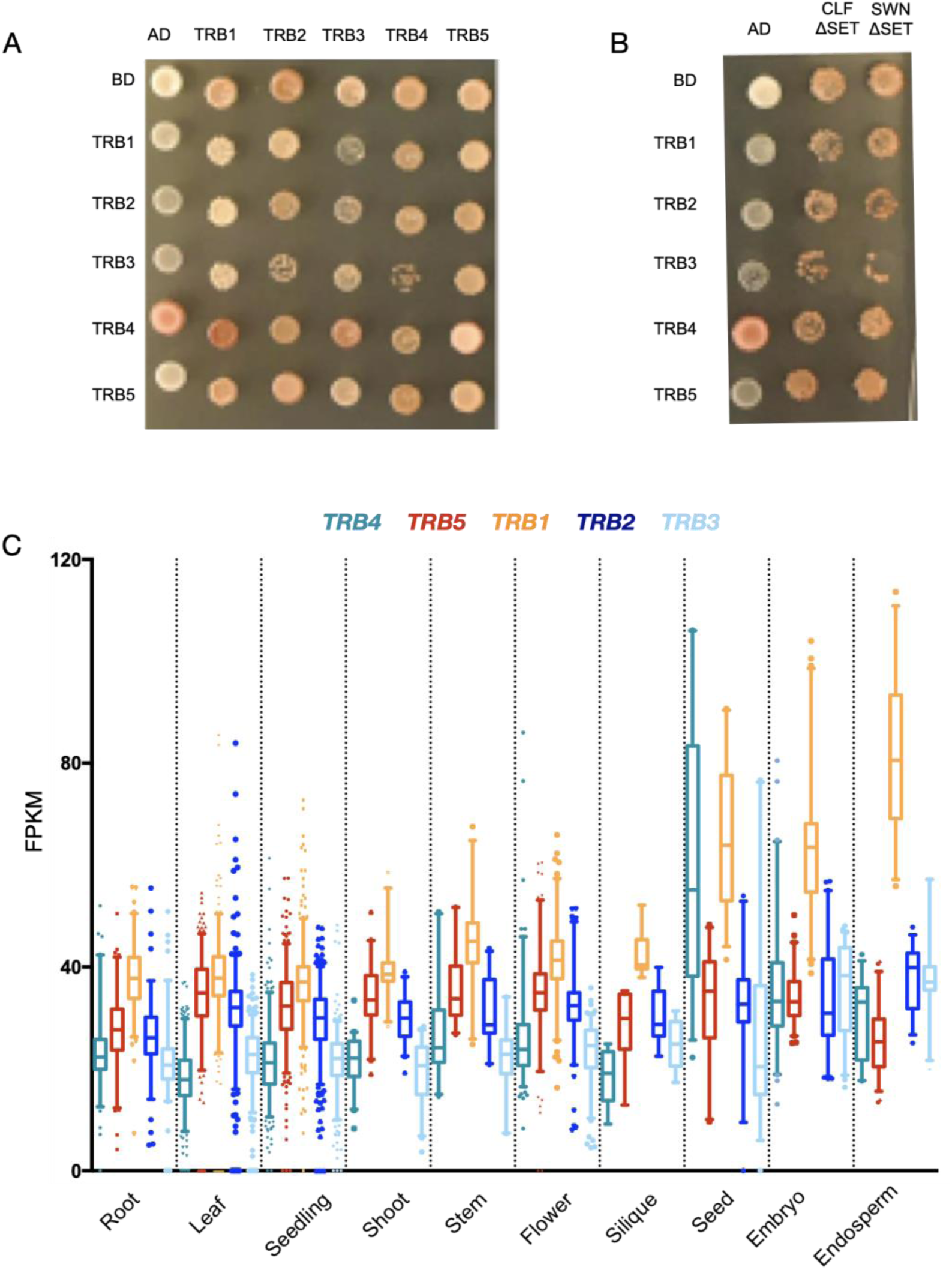
Supplemental information related to Figure 3. **(A, B)** Growth of zygotes on synthetic medium lacking leucine and tryptophan, selecting for the presence of the bait and prey vectors for interactions scored in Figure 3A (**A**) and Figure 6B **(B). (C)** Expression level of *TRB1, TRB2, TRB3, TRB4* and *TRB5* in different Arabidopsis tissues issued from available RNA-seq datasets (data extracted from Arabidopsis RNA-seq Database - http://ipf.sustech.edu.cn/pub/athrna/). All 5 genes are ubiquitously expressed to similar levels, except in the embryo and endosperm that show higher *TRB1* transcript levels.

**Supplementary Figure 4:**
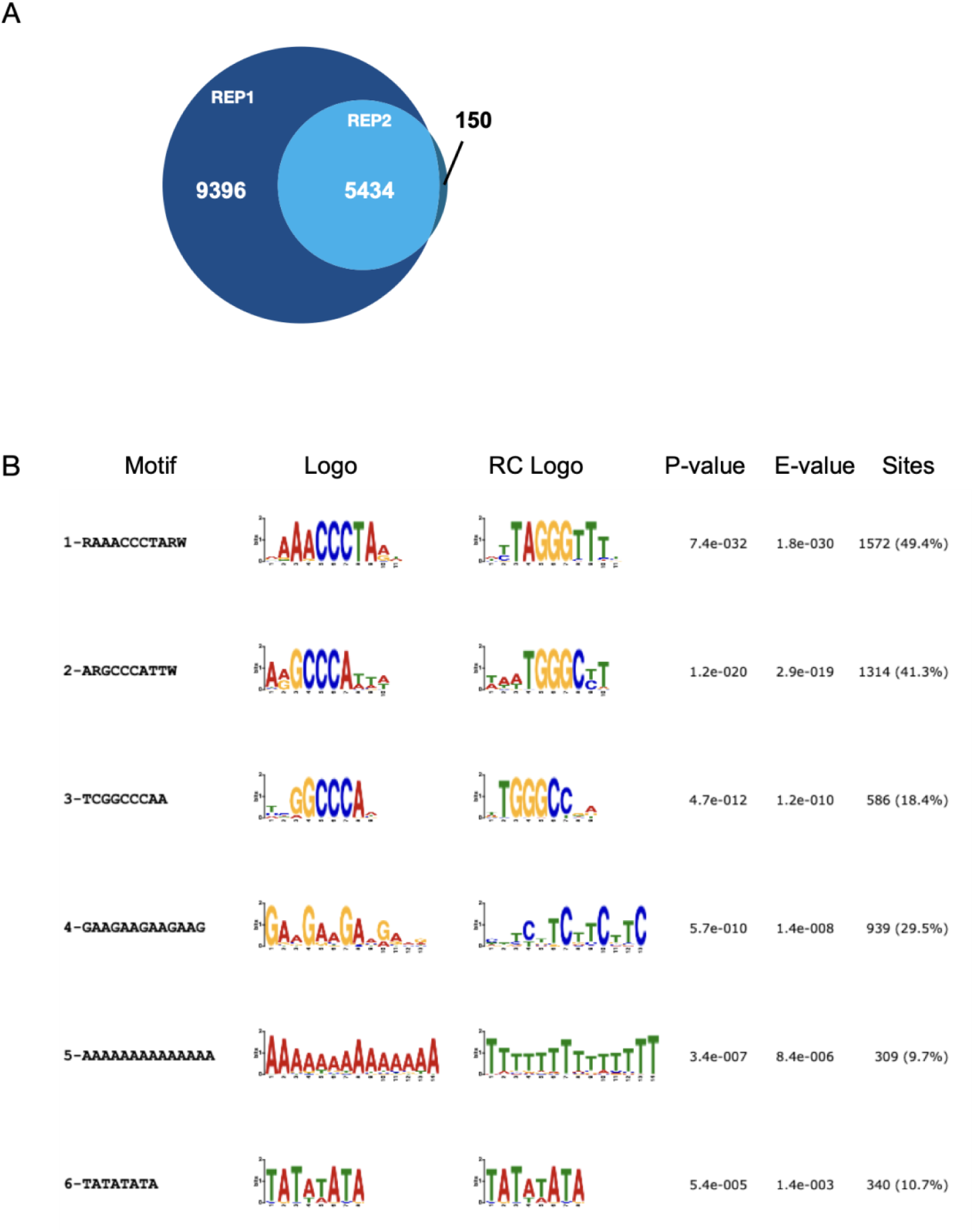
Supplemental information related to Figure 4. **(A)** Venn diagram showing overlap between the identified TRB4 targets in the two biological replicates. **(B)** The six most abundant DNA sequence motifs identified by MEME at TRB4 binding sites. Motif 1 corresponds to the telobox, motifs 2 and 3 corresponds to ‘site II motif’ (*TGGGCY*) (Gaspin et al., 2010).

**Supplementary Figure 5:**
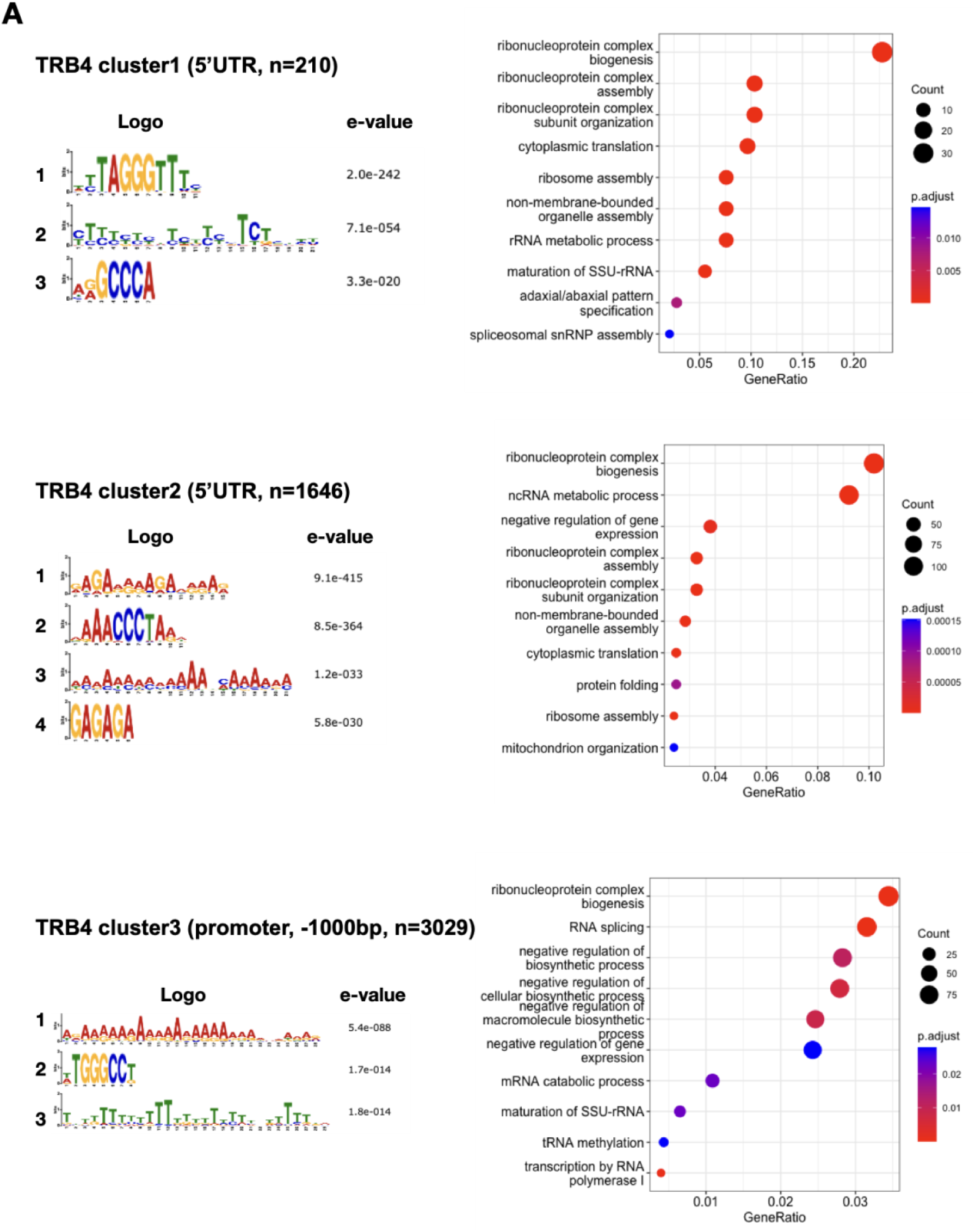
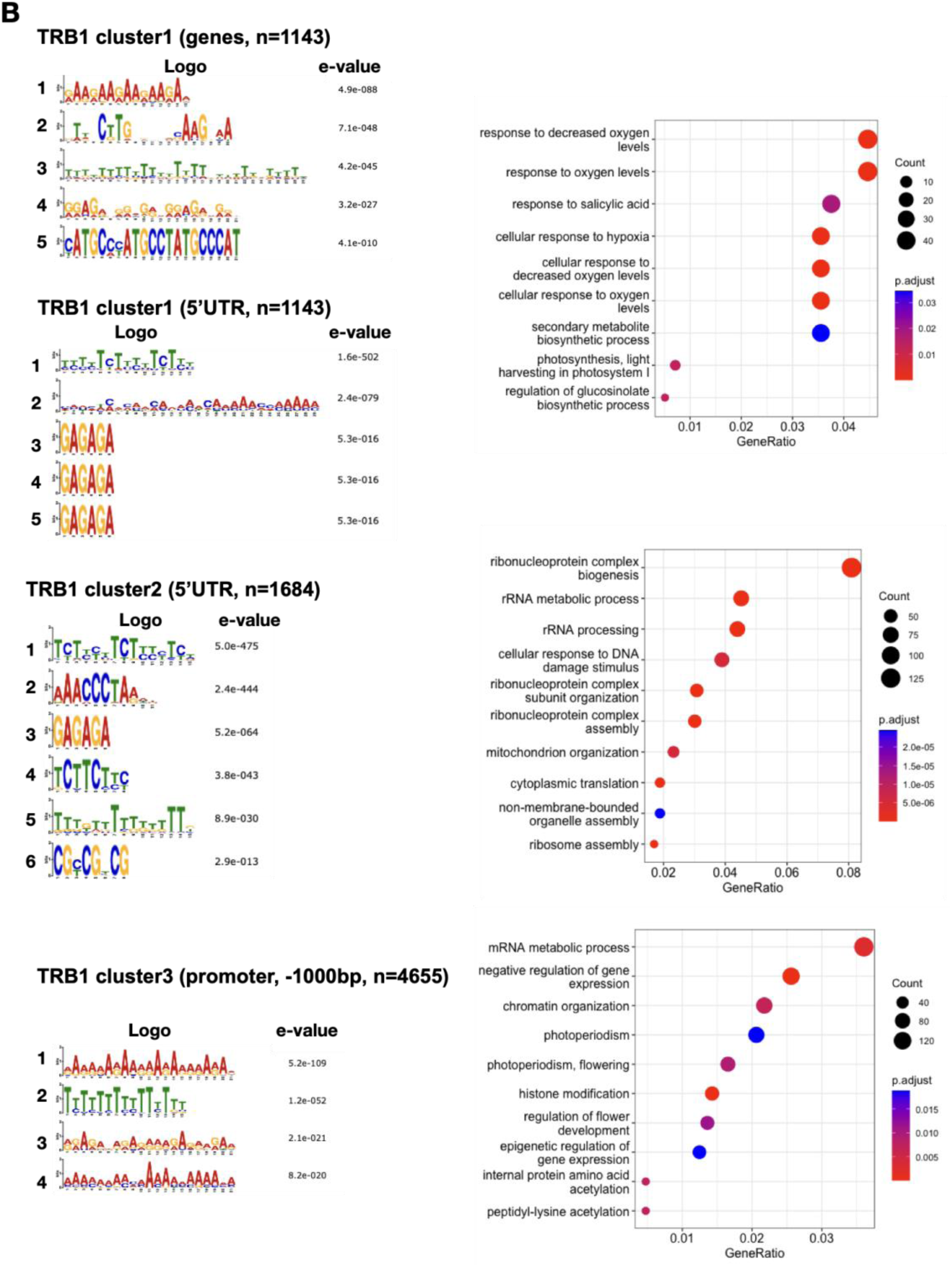
Supplemental information related to Figure 5. (**A, B**) Left: MEME predictions of up to 6 DNA sequence motifs within the 5’UTR, the promoters (-1000bp from the TTS) of the TRB4 (**A**) or TRB1 (**B**) target genes in the three clusters defined in Figure 5A-B. Right: GO-term enrichment of genes corresponding to the three cluster of TRB4 (**A**) and TRB1 (**B**) target genes defined in Figure 5A-B.

**Supplementary Figure 6:**
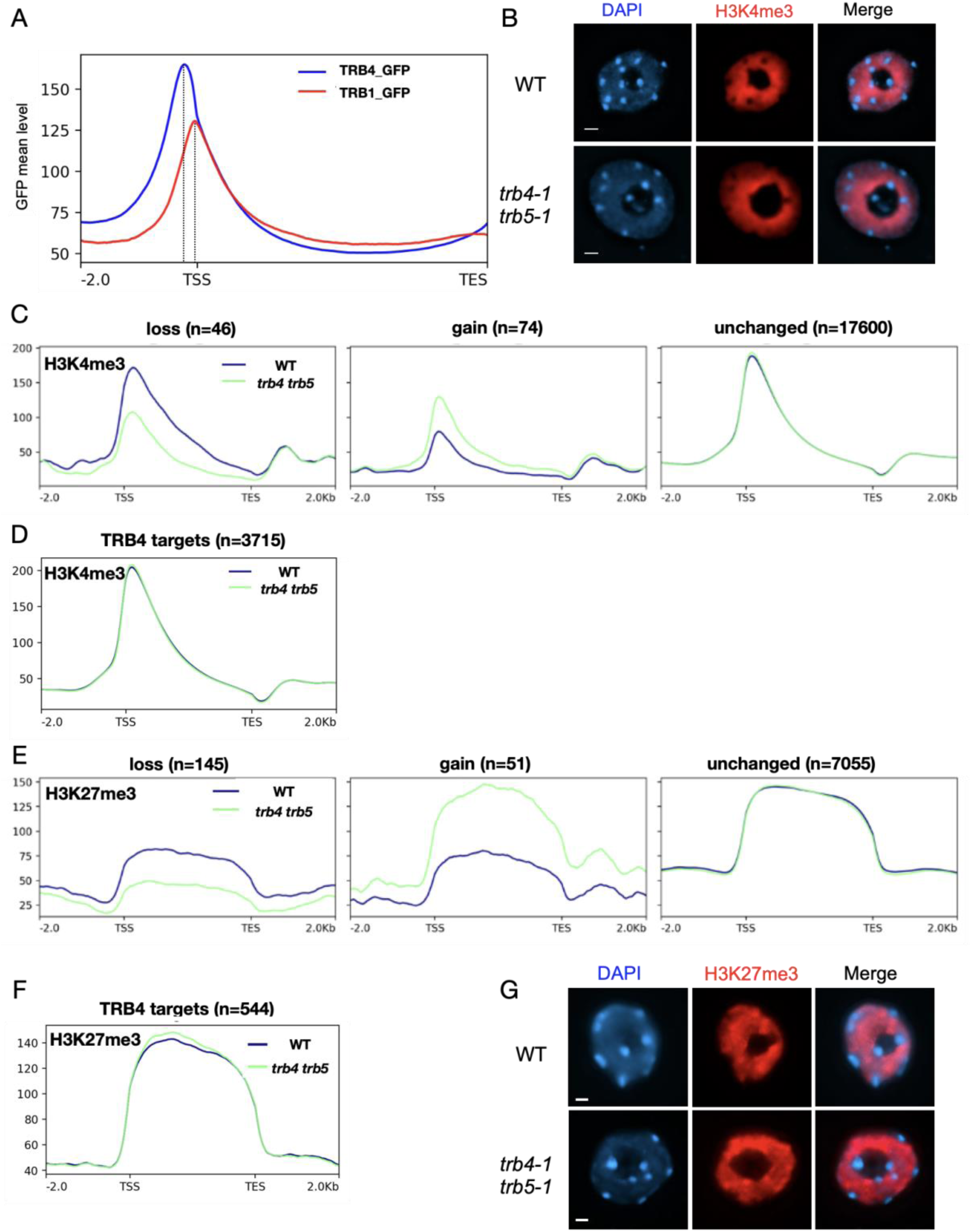
Supplemental information related to Figure 6. **(A)** Metagene plot showing ChIP-seq signals of TRB4-GFP or TRB1-GFP over genes enriched in H3K4me3. TRB4 peaks upstream of TRB1. **(B)** Representative mesophyll leaf nuclei from wild type and *trb4-1 trb5-1* mutant plants, immunostained for H3K4me3. Maximum projections are shown. Scale presents 1 μm. **(C, E)** Metagene plot presentations of H3K4me3 **(C)** and H3K27me3 **(E)** enrichment along genes and 2 kb up and downstream of TSS and TTS with loss, gain or unchanged levels of the histone modifications in *trb4-1 trb5-1* mutants. The number of genes presented in each graph is indicated on the top. **(D, F)** Metagene plot showing enrichment of H3K4me3 **(D)** and H3K27me3 **(F)** over TRB4-target genes. (**G**) Representative mesophyll leaf nuclei from wild type and *trb4-1 trb5-1* mutant plants, immunostained for H3K27me3. Maximum projections are shown. Scale presents 1 μm.

